# Single-cell transcriptomics of the *Drosophila* wing disc reveals instructive epithelium-to-myoblast interactions

**DOI:** 10.1101/2020.07.27.222976

**Authors:** Nicholas J. Everetts, Melanie I. Worley, Riku Yasutomi, Nir Yosef, Iswar K. Hariharan

**Author notes:** co-first authorship (Nicholas J. Everetts, Melanie I. Worley).

## Abstract

In both vertebrates and invertebrates, generating a functional appendage requires interactions between ectoderm-derived epithelia and mesoderm-derived cells. To investigate such interactions, we used single-cell transcriptomics to generate a cell atlas of the *Drosophila* wing disc at two time points during development. Using these data, we investigate gene expression using a multi-layered model of the wing disc and catalogued ligand-receptor pairs that could mediate signaling between epithelial cells and adult muscle precursors (AMPs). We found that localized expression of the FGF ligands, Thisbe and Pyramus, in the disc epithelium regulates the number and location of the AMPs. In addition, Hedgehog ligand from the epithelium activates a specific transcriptional program within adjacent AMP cells, which is critical for proper formation of a subset of the direct flight muscles. More generally, our annotated atlas provides a global view of potential cell-cell interactions between subpopulations of epithelial and myogenic cells.

## Introduction

The development of multicellular eukaryotes gives rise to organs that are composed of cells of many types, typically derived from different germ layers such as the ectoderm and the mesoderm. There is increasing evidence that interactions between different cell types during development play a role in ensuring that the appropriate quantity of cells of each type is present in the fully formed adult organ (Ribatti & Santoiemma, 2014). A particularly well-studied example of such heterotypic interactions occurs during the development of the vertebrate limb, where signals are exchanged between the apical ectodermal ridge and the underlying mesoderm (Delgado & Torres, 2017). Additionally, aberrant heterotypic interactions have been observed in many pathological conditions such as cancer (Werb & Lu, 2015). For example, interactions between breast cancer cells and myofibroblasts are thought to play an important role in tumor progression (Orimo & Weinberg, 2006).

While vertebrate limbs are relatively complex structures, the *Drosophila* wing-imaginal disc, the larval primordium of the adult wing and thorax, is ideally suited to the study of cell-cell interactions in the context of organ development because of its relative simplicity and amenability to genetic analysis (Waddington, 1940; Cohen, 1993; Neto-Silva et al., 2009). The wing-imaginal disc is composed of epithelial cells that form a sac-like structure (comprised of the columnar cells of the disc proper and the squamous cells of the peripodial epithelium) and a population of adult muscle precursors (AMPs) that resides between the epithelial cells of the disc proper and the underlying basement membrane (**Figure 1A**). The epithelial portion of the disc derives from a primordium of approximately 30 cells from the embryonic ectoderm that are specified during embryogenesis (Mandaravally Madhavan & Schneiderman, 1977; Worley et al., 2013; Requena et al., 2017). The AMPs, originally referred to as adepithelial cells (Poodry & Schneiderman, 1970), represent a subset of cells from the embryonic mesoderm that generate the adult flight muscles (Bate et al., 1991; Fernandes et al., 1991). The AMPs underlie the dorsal portion of the wing disc, the notum, which is the primordium of the dorsal thorax. During metamorphosis, these AMPs generate multiple muscle fibers which comprise the direct and indirect flight muscles (DFMs and IFMs, respectively) (reviewed by Bothe & Baylies, 2016; Gunage et al., 2017; Laurichesse & Soler, 2020).

**Figure 1.**
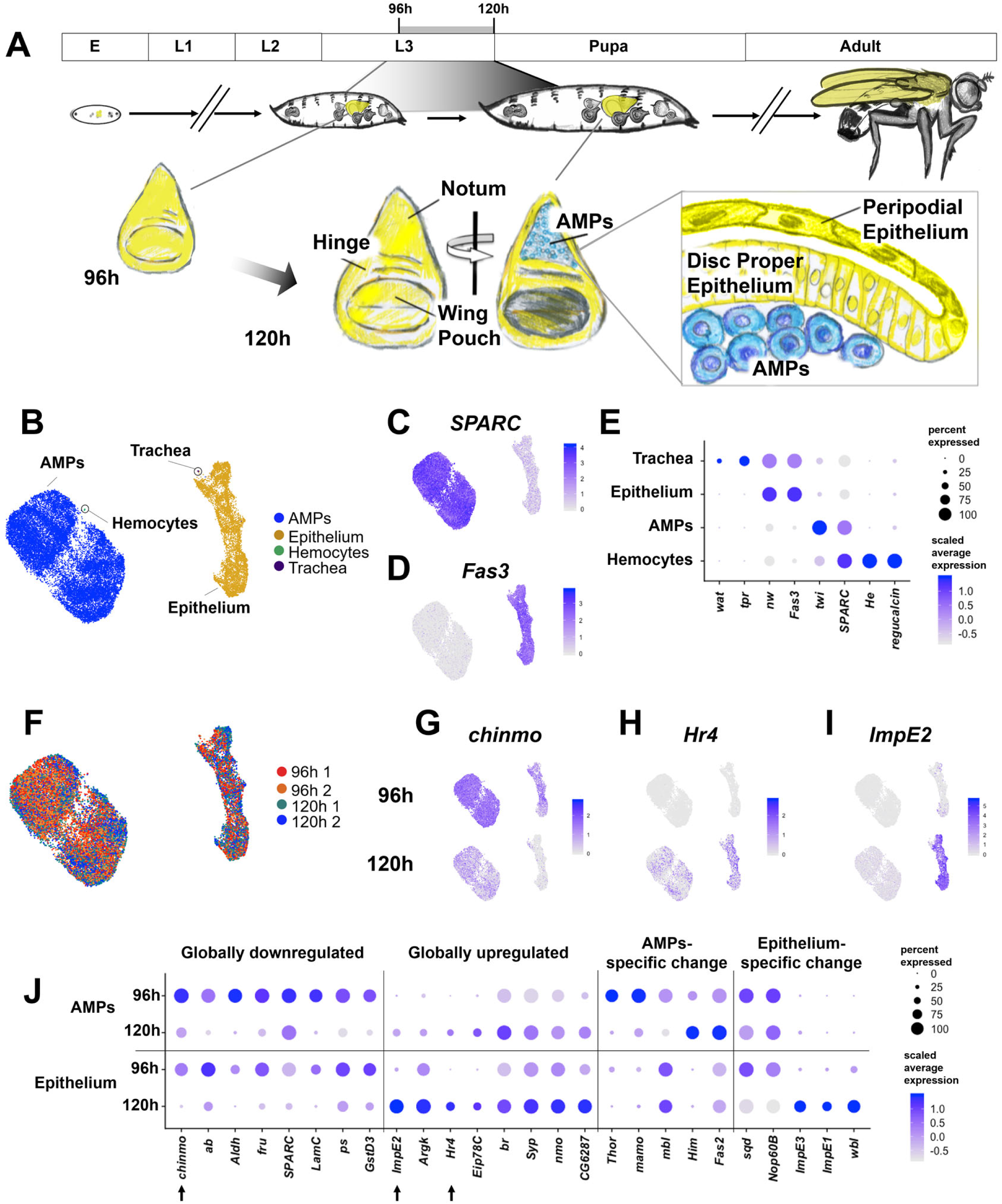
Cell atlas of the developing wing-imaginal disc. (**A**) Timeline of *Drosophila* development: embryo (E), larval phases (L1-L3), pupa, and adult. Diagram of the wing-imaginal disc within the 3rd larval instar (L3) from mid (96h AEL) and late (120h AEL) time points. The epithelial cells of the wing disc become the adult wing blade, hinge, and the majority of the dorsal thorax (shown in yellow). The myoblasts associated with the tissue are the adult muscle precursors (AMPs) and are shown in blue. The AMPs reside on the basal surface of the disc proper epithelium. During metamorphosis the AMPs undergo cell fusion events to generate the adult flight muscles within the thorax. (**B**) UMAP of harmonized single-cell datasets (two biological replicates each of 96h and 120h AEL), colored by identified cell types (AMPs, epithelium, hemocytes, and trachea). The AMPs and epithelial cells can be distinguished by high expression levels of *SPARC* (**C**) and *Fas3* (**D**) respectively, as shown on the UMAP. Expression scale bars correspond to normalized transcript counts (scaled by total UMIs per cell) on a natural-log scale. (**E**) Dot plot summarizing gene expression for known markers of each of the identified cell types. For a gene of interest, the size of the dot indicates the percent of cells that express the gene and the color of the dot indicates the relative gene expression level (see **Materials and Methods**). (**F)** Cells of the UMAP colored by batch (two biological replicates each for 96h and 120h AEL). **(G-I)** Expression (visualized via UMAP) for *chinmo* (**G**)*, Hr4* (**H**), and *ImpE2* (**I**), which are differentially-expressed between the two time points (false discovery rate [FDR] < 0.05 in all pairwise comparisons, see **Materials and Methods**). UMAPs are separated by time points (96h and 120h) to illustrate changing expression levels of these genes. Color scales for UMAPs correspond to normalized (by total UMI) counts on a natural-log scale. (**J**) Dot plot summarizing gene expression changes from 96h to 120h, that were observed within both the epithelium and AMPs (“Globally downregulated / upregulated”) or mostly within a single cell type (“AMP-specific / Epithelium-specific changes”). Genes shown are differentially-expressed between the two time points in the indicated populations (FDR < 0.05 for all pairwise comparisons). Arrows highlight the genes shown on the UMAPs in **G-I**. For visualization on the dot plot, normalized expression counts for each gene are standardized to have mean of 0 and a standard deviation of 1.

The mechanisms that influence a seemingly uniform population of AMPs to generate different types of flight muscles each composed of multiple distinct fibers are not known. The AMPs are generated by an asymmetric division of a muscle founder cell during embryogenesis; one daughter cell becomes an AMP while the other generates precursors of larval muscles (Bate *et al*., 1991). In the second thoracic segment, dorsal clusters of AMPs, which express a segment-specific combination of Hox genes (Roy & VijayRaghavan, 1997), become associated with the wing disc, remain in the notum region, and proliferate via symmetric cell divisions. At the onset of the third larval instar (L3), the protein Wingless (Wg; a ligand of the Wnt signaling pathway) secreted from epithelial cells of the notum triggers a switch within the AMPs to an asymmetric pattern of cell division (Gunage et al., 2014). The precursors of the indirect and direct flight muscles can be distinguished by higher levels of expression of the transcriptional regulators Vestigial or Cut (Sudarsan et al., 2001). The elevated Vestigial expression in the IFM precursors is maintained by expression of Wg ligand in the notum (Sudarsan et al., 2001).

Two main issues central to the mechanisms that regulate proliferation and cell-fate specification in the AMPs, however, remain largely unanswered. First, it has been suggested that the notum acts as a “dynamic niche” that both regulates the survival of AMPs and guides their specification (Gunage et al., 2017). However, the signals emanating from the epithelial cells to either regulate AMP numbers or maintain them in the notum region of the wing disc have not been identified. Second, it has always been assumed that extrinsic signals, likely from the disc epithelium, function during the larval stage to direct subsets of the AMPs to become precursors of specific types of muscles (Gunage et al., 2017). However, with the exception of Wg, such signals remain unidentified.

Single-cell transcriptomics provides a powerful paradigm for mapping the cellular composition of developing organs (Schier, 2020), including vertebrate appendages (Fabre et al., 2018; Cao et al., 2019; Feregrino et al., 2019), and has been successfully utilized for characterizing the *Drosophila* wing disc at single snapshots during development (Deng et al., 2019; Bageritz et al., 2019; Zappia et al., 2019). Beyond cataloging cell types, transcriptome-scale analysis of single cells opens the way for a comprehensive evaluation of how interactions between cells may facilitate development (Satija et al., 2015; Karaiskos et al., 2017; Deng et al., 2019; Bageritz et al., 2019). Analysis of cell-cell interactions in this context can benefit greatly from information about the physical organization of the tissue in question, and particularly the position of different cell types in the tissue. Since in most prevalent protocols this information is lost, computational methods are used to infer it, usually based on the expression of landmark genes (Satija et al., 2015; Karaiskos et al., 2017; Deng et al., 2019; Bageritz et al., 2019). Complementary to spatial localization are approaches that have examined the expression of receptors, ligands, and downstream molecules to predict which cell subsets interact and by what mechanisms (Vento-Tormo et al., 2018; Browaeys et al., 2020).

The combination of these approaches, namely estimating the physical context of each cell and mapping the expression of extracellular cues such as receptor-ligand pairs, can provide a powerful tool for studying organ development. Here, we couple these two approaches to identify heterotypic interactions that are crucial for disc development, focusing on signaling between the disc epithelium and the AMPs. To this end, we collected single cell RNA-sequencing (scRNAseq) data from two developmental time points and derived a comprehensive view of cell subsets, their spatial organization, and cell-cell communications. We identified a set of developmentally-regulated transcriptional changes in each of these populations, some global and others restricted to specific spatial domains. We obtained a comprehensive view of potential cell-cell interactions by generating a transcriptional model of the wing disc and examining the expression of receptors and ligands. From our analysis, we identified key interactions between the disc epithelium and the adepithelial AMPs. Specifically, we show that FGF ligands emanating from the disc epithelium create an AMP niche that regulates AMP numbers and restricts them to the region of the notum. Furthermore, we find that Hedgehog ligand from the disc epithelium specifies aspects of cell fate in a subset of AMPs and provide evidence for stereotyped AMP movement during development. Beyond these examples, our annotated dataset provides a resource for spatiotemporal cellular composition in the developing wing disc and points to additional potential heterotypic interactions between epithelial cells and AMPs.

## Results

### Single-cell transcriptomics of the developing wing disc identifies transcriptional changes in the epithelium and the AMPs

The wing disc is composed of multiple cell types, including the columnar cells of the disc proper, the squamous cells of the peripodial epithelium, and the mesoderm-derived AMPs (**Figure 1A**). In addition, the wing disc is in intimate contact with branches of the tracheal system and circulating blood cells called hemocytes. With the goal of generating a spatiotemporal atlas of the developing wing disc, we used single-cell RNA sequencing to collect transcriptional profiles of cells at mid and late 3rd instar, which correspond to 96h and 120h after egg lay (AEL) (**Figure 1A**). Two biological replicates were obtained at each time point that, after filtering for low-quality cells, generated data from 6,922 and 7,091 cells in the 96h samples and 7,453 and 5,550 cells in the 120h samples. Harmonization of the different samples and dimensionality reduction (which provides the basis for visualization and clustering) was performed using scVI (Lopez et al., 2018). Clustering and differential expression analysis was done with the Seurat v3 R package (Stuart et al., 2019), and two-dimensional visualization of the data was performed with UMAP (McInnes et al., 2018) (see **Materials and Methods**).

Our single-cell analysis identified four major cell types via known gene markers (**Figure 1B**). The largest cell type recovered was identified as the AMPs by high transcript levels for the collagen-binding protein *SPARC* (*Secreted protein, acidic, cysteine-rich*) (Butler et al., 2003) and mesodermal transcription factor *twist* (*twi*) (Bate et al., 1991) (**Figure 1C, E**). The wing disc epithelial cells (disc proper and peripodial epithelium) were identified by high expression levels of the *Fasciclin 3* (*Fas3*), which encodes for a cell adhesion molecule that marks thoracic imaginal discs (Bate & Martinez Arias, 1991), and the gene *narrow* (*nw*), which is known to regulate wing-disc size (Lindsley & Zimm, 1992) (**Figure 1D-E**). The few tracheal cells were identified by high expression of the genes *waterproof* (*wat*) and *tracheal-prostasin* (*tpr*) (Jaspers et al., 2014; Drees et al., 2019), and the hemocytes were distinguished by high expression levels of *Hemese* (*He*) and *regucalcin* (Kurucz et al., 2003; Klebes et al., 2005) (**Figure 1E, Figure 1 - figure supplement 1**). We observed a similar set of marker genes as other recently published wing disc scRNAseq analyses (Deng et al., 2019; Bageritz et al., 2019; Zappia et al., 2019). Altogether, we recovered profiles for 19,885 adult muscle precursors (AMPs), 7,104 wing disc epithelial cells, 15 tracheal cells, and 12 hemocytes. These major cell types were captured at both developmental time points (**Figure 1F**). Notably, our dataset shows a dramatic overrepresentation of AMPs with an observed ratio of around 2.8 AMPs to 1 epithelial cell instead of the expected 1:12, based on the estimate of the wing disc being comprised of 2,500 AMPs (Gunage et al., 2014) and 30,000 epithelium cells at the end of the 3rd larval instar (Martín et al., 2009). The enrichment is likely the result of our collagenase-based dissociation protocol (see **Materials and Methods**) that appears to have dissociated AMPs far more effectively than epithelial cells. Thus, the enrichment of AMPs has enabled an especially detailed analysis of this cell type, which has previously received less attention.

In the developmental window between 96h and 120h, the size of the tissue more than quadruples in size due to cell growth and proliferation (Steiner, 1976; McClure & Schubiger, 2005). During that process, particular cell types are further specified, such as the cells of the future wing margin within the disc pouch. Changes in expression levels during this developmental window have been previously reported for many genes (Li & White, 2003), but expression changes in epithelial cells and AMPs have not been evaluated separately. We examined gene expression changes within these two major cell types between 96h and 120h and classified the changes as being organ-wide or cell type specific (false discovery rate [FDR] < 0.05 in all pairwise comparisons; see **Methods and Methods** for details of our differential expression analysis). The gene *Chronologically inappropriate morphogenesis* (*chinmo*), which encodes for a BTB-zinc finger transcription factor and is known to be downregulated during development (Narbonne-Reveau & Maurange, 2019), was observed to be downregulated in both major cell types (**Figure 1G, 1J**). Conversely, genes induced by ecdysone, such as *Hormone receptor 4* (*Hr4*), *Ecdysone-inducible gene E2* (*ImpE2*) (Osterbur et al., 1988), and *Arginine kinase* (*ArgK*) (James & Collier, 1992), were upregulated in both the epithelium and AMPs (**Figure 1H-J**). Interestingly, *ImpE2* was more strongly induced within the epithelium than the AMPs, a trend that was even more dramatically observed for two additional ecdysone-inducible genes, *ImpE1* and *ImpE3* (**Figure 1J**). These results suggest that the epithelium and AMPs may respond differently to ecdysone expression - a critical hormonal change that leads to termination of the growth phase and starting of metamorphosis (**Figure 1A**).

It is interesting to note that some genes stayed consistently expressed in one cell type while being dynamic in another. For example, *muscleblind* (*mbl*) (Begemann et al., 1997), which encodes for a RNA binding protein that regulates alternative splicing (Ho et al., 2004), was specifically downregulated in the AMPs but remained constantly expressed in the epithelium (**Figure 1J**), suggesting a mechanism where ecdysone could alter splice-site choice within specific cell types. A complete list of the genes that exhibit differential expression over time in either the epithelium and/or the AMPs is provided in **Supplementary file 1**.

### Major transcriptional differences between epithelial cells reflect their proximodistal position

To define the signals that might be exchanged between the epithelium and the AMPs, we first characterized the cell types within each of these populations separately. The wing disc epithelium is often divided into four broad domains of the notum, hinge, pouch, and peripodial epithelium based both on morphology and gene expression patterns. Although the exact boundaries between these domains are somewhat ambiguous, the genes encoding the transcription factors *nubbin* (*nub*) and *Zn finger homeodomain 2* (*zfh2*) are often used to define the pouch and hinge, respectively (Zirin & Mann, 2007; Terriente et al., 2008; Ayala-Camargo et al., 2013), and are shown for wing discs at 96h and 120h AEL (**Figure 2A, B**). Based on their contributions to adult structures, the three domains of the disc proper (notum, hinge, and pouch) define the proximodistal axis of the wing disc, with the notum being the most proximal and the pouch being the most distal. We visualized the expression of *nub* and *zfh2* within our data (**Figure 2C, D**), as well as marker genes for the anterior notum, *eyegone* (*eyg*) (**Figure 2E**) and for the squamous portion of the peripodial epithelium *Ultrabithorax* (*Ubx*) (**Figure 2F**). This analysis revealed localized expression of these marker genes as visualized by UMAP. These results therefore suggest that the cells are largely grouped by their putative region in the tissue and that the proximodistal axis is a primary source of variation in the data.

**Figure 2.**
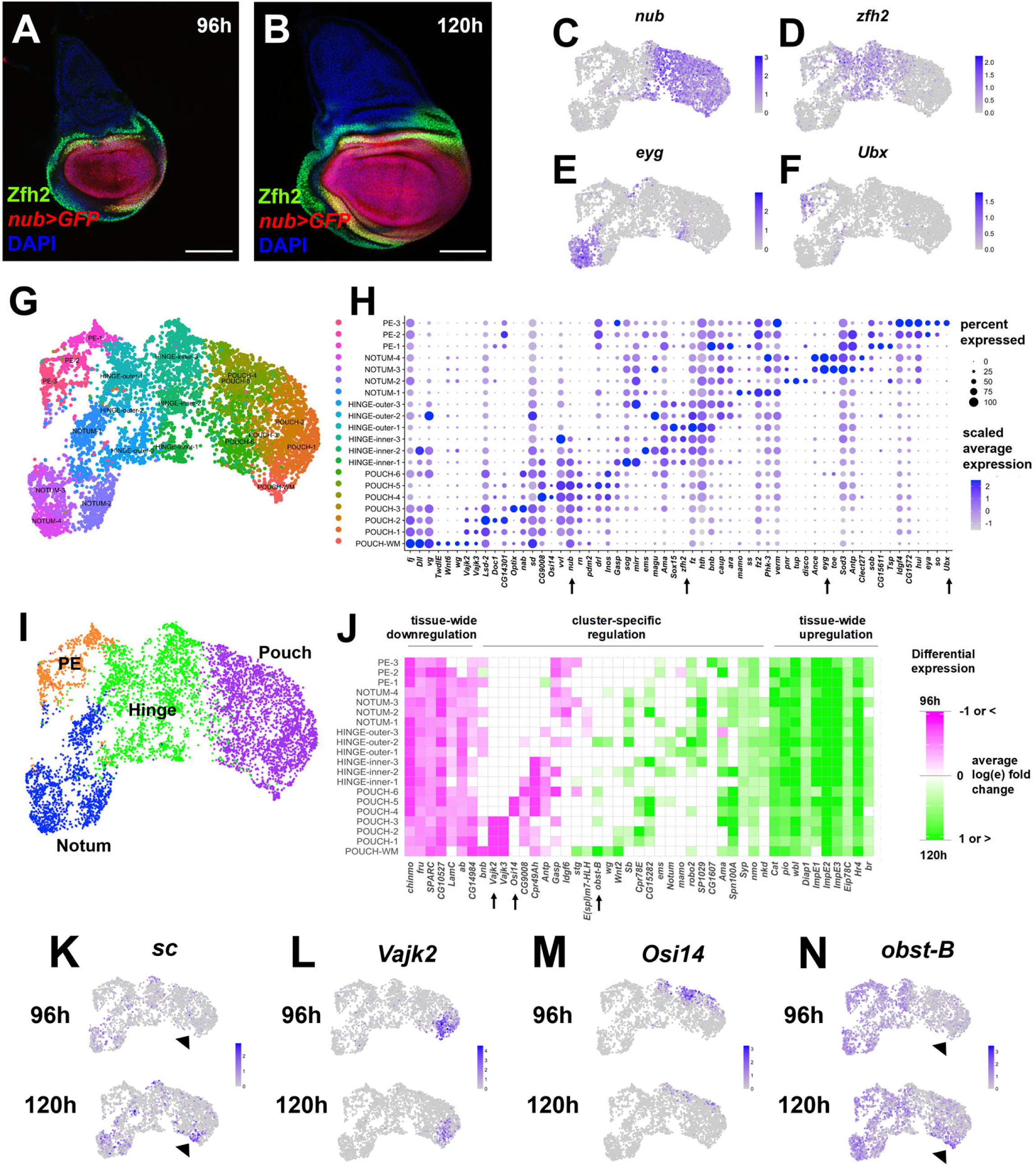
Temporal cell atlas of the wing disc epithelium. (**A, B**) Mid (**A**) and late 3^rd^ instar (**B**) wing-imaginal discs with *nub-GAL4* driving the expression of GFP (red) to mark the pouch and anti-Zfh2 (green) to mark the hinge. Wing discs are oriented such that anterior is left, posterior is right. Images are max projections across all image slices. Note that the wing disc has a similar domain identity at mid and late 3rd instar. (**C-F**) UMAPs of the harmonized epithelium cells, with cells colored by their expression levels of *nub* (**C**)*, zfh2* (**D**)*, eyg* (**E**), and *Ubx* (**F**). Note that cells with high expression of each of these genes are largely grouped together on the UMAP. (**G**) UMAP of epithelium cells, colored by cluster identities as determined by Seurat. Cluster names were manually provided based on identity determined by marker gene expression. Note the relative organization of the UMAP matches that of the proximal-distal axis of the wing disc (also note the cluster identification from cell mapping to virtual tissue model in **Figure 4**). (**H**) Dot plot showing the level and percent of differentially-expressed (FDR < 0.05) marker gene expression for the cell clusters. Note that there are many genes that are expressed in a gradient. Arrows indicate the genes with highlighted expression levels on the UMAPs above. (**I**) UMAP of epithelium cells, where cell clusters have been merged into the major domains (pouch, hinge, notum, and peripodial epithelium (PE)). (**J**) Plot showing differential gene expression for developmentally-regulated genes for the epithelium cell clusters. All genes shown were differentially-expressed between time points in at least once cluster (FDR < 0.05 for all pairwise comparisons, see **Materials and Methods**). Non-significant fold-changes were set to have a value of zero within the plot. Natural-log fold-change of expression was calculated for each of the cell clusters between cells of 96h and 120h and is capped at +/− 1 for better visualization. Genes were selected to highlight different expression dynamics, as either changing throughout most of the clusters (“tissue-wide downregulation / upregulation”) or only having cluster-specific changes (“cluster-specific regulation”). Arrows indicate the genes with expression levels shown on the UMAPs below. (**K-N**) UMAPs of disc epithelium cells separated by time points (96h and 120h), with cells colored by gene expression as labeled. (**K**) The pro-neural gene *scute* (*sc*) is a marker of the wing margin (indicated by the arrowheads) and is expressed at higher levels in the 120h sample. While *sc* does not pass our differential-expression criteria between time points (see **Materials and Methods**), it should be noted that more than 75% of cells within the margin cluster originate from the 120h time point (see **Figure 2 - figure supplement 2**). (**L**) *Vajk2* and (**L**) *Osiris 14* (*Osi14*) are differentially-expressed at higher levels at 96h within specific regions of the pouch. (**N**) *obstructor-B* (*obst-B*) is specifically upregulated within the wing margin (“POUCH-WM”), indicated on the UMAP by the arrowheads, while expression in many other clusters remains relatively unchanged. Color scales for UMAPs correspond to normalized (by total UMI) counts on a natural-log scale. Microscopy scale bars = 100 μm.

Interestingly, while the proximodistal axis was a primary feature in stratifying cells within our analysis, the anteroposterior axis stratified the data to a lesser degree. The cells that generate the anterior compartment of the disc arise from embryonic subpopulations distinct from those that generate the posterior, and the two compartments remain physically separate throughout development (Garcia-Bellido et al., 1973; Mandaravally Madhavan & Schneiderman, 1977; Worley et al., 2013; Requena et al., 2017). We classified cells as originating from either the anterior or posterior compartment based on marker genes, such as transcriptional activator *cubitus interruptus* (*ci*) that is expressed within anterior compartment (**Figure 2 - figure supplement 1A-E**). We found that the concise representation of the data in two dimensions (with UMAP) resulted in stratification of the cells based on the proximodistal axis first, with secondary stratification of cells within each tissue region into anterior or posterior. Furthermore, we found that comparison of cells across the proximodistal axis results in more differentially-expressed genes, compared with the anteroposterior axis (**Figure 2 - figure supplement 1F**). Thus, the cells at the same proximodistal position but in different anteroposterior compartments tend to be more similar transcriptionally, even though they diverged early in development (latest common ancestor at least 10 cell divisions ago; Garcia-Bellido et al., 1973).

To further characterize the epithelial compartment, we stratified the cells into 20 groups using unsupervised clustering (**Figure 2G**) and found genes that marked each cluster using differential expression analysis (see **Materials and Methods**). While the clusters can be efficiently classified as originating from the notum, hinge, pouch, and peripodial epithelium (**Figure 2I**), we used the differential expression analysis in conjunction with known marker genes to provide further stratification to subsets of cells within each region (**Figure 2H, I;** see also **Figure 4 - figure supplement 1**). For example, while we classify several clusters as originating from the hinge due to high expression of known markers *zfh2* and *Sox box protein 15* (*Sox15*) (Terriente et al., 2008; Miller et al., 2009), we further stratify these clusters as being inner and outer hinge based on the expression of genes like *ventral veins lacking* (*vvl*; also known as *drifter*) and *nub*, both of which have strong expression within and around the pouch (Certel et al., 2000; Zirin & Mann, 2007).

We next used the clustering analysis to explore transcriptional changes during development that may be restricted to specific regions of the wing disc epithelium. In total, we found that 337 genes were significantly downregulated and 408 genes were significantly upregulated in one or more clusters of epithelial cells (for details on differential expression significance, see **Materials and Methods**). Examples of different types of gene regulatory events that occurred within the epithelial cells over this time, which include tissue-wide and cluster-specific are shown in **Figure 2J**. We noted the increased expression of the pro-neural gene *scute* (*sc*) within the wing margin (**Figure 2K**), which coincides with the specification of neurons in the future wing margin (Romani et al., 1989). In addition, we observed region-specific downregulation of *Vajk2* and *Osiris 14* (*Osi14*) within the inner and outer pouch clusters (**Figure 2J, 2M, 2L**). Both of these genes are expressed strongly at 96h and have reduced expression at 120h. In contrast, our analysis also identified genes that are expressed more widely but only upregulated in a cluster-specific manner. One such example is *obstructor-B* (*obst-B*) (**Figure 2N**), which is widely expressed but specifically upregulated in the wing margin. Interestingly, not all genes changed expression in the same direction over the course of development. We found 19 genes that were significantly upregulated within some clusters while simultaneously being significantly downregulated within others. One example is *string* (*stg*) (Edgar & O’Farrell, 1990), which encodes a regulator of the cell cycle and is upregulated within the wing margin while being downregulated in other regions of the disc (**Figure 2J**). Thus, even though the major cell types are established by 96h of development (**Figure 2A, B; Figure 2 - figure supplement 2**), we still find evidence of further pattern refinement via highly-localized gene expression changes. The list of the genes that exhibit differential expression within clusters between the two time points is provided in **Supplementary file 2**.

### Cell-type identities among the AMPs are consolidated much later than in the epithelium

We next analyzed cells categorized as AMPs (**Figure 3A**). Initial analysis of these data showed a clear partition of the cells with respect to two primary features: cell cycle phase and the sex of the donor fly (**Figure 3 - figure supplements 1 and 2**). We utilized scVI to recompute a low-dimensional representation of the cells (later used for clustering and visualization) while suppressing the effects of these covariates, and thus obtained a clearer view of other aspects of AMP cell biology (see **Materials and Methods**).

**Figure 3.**
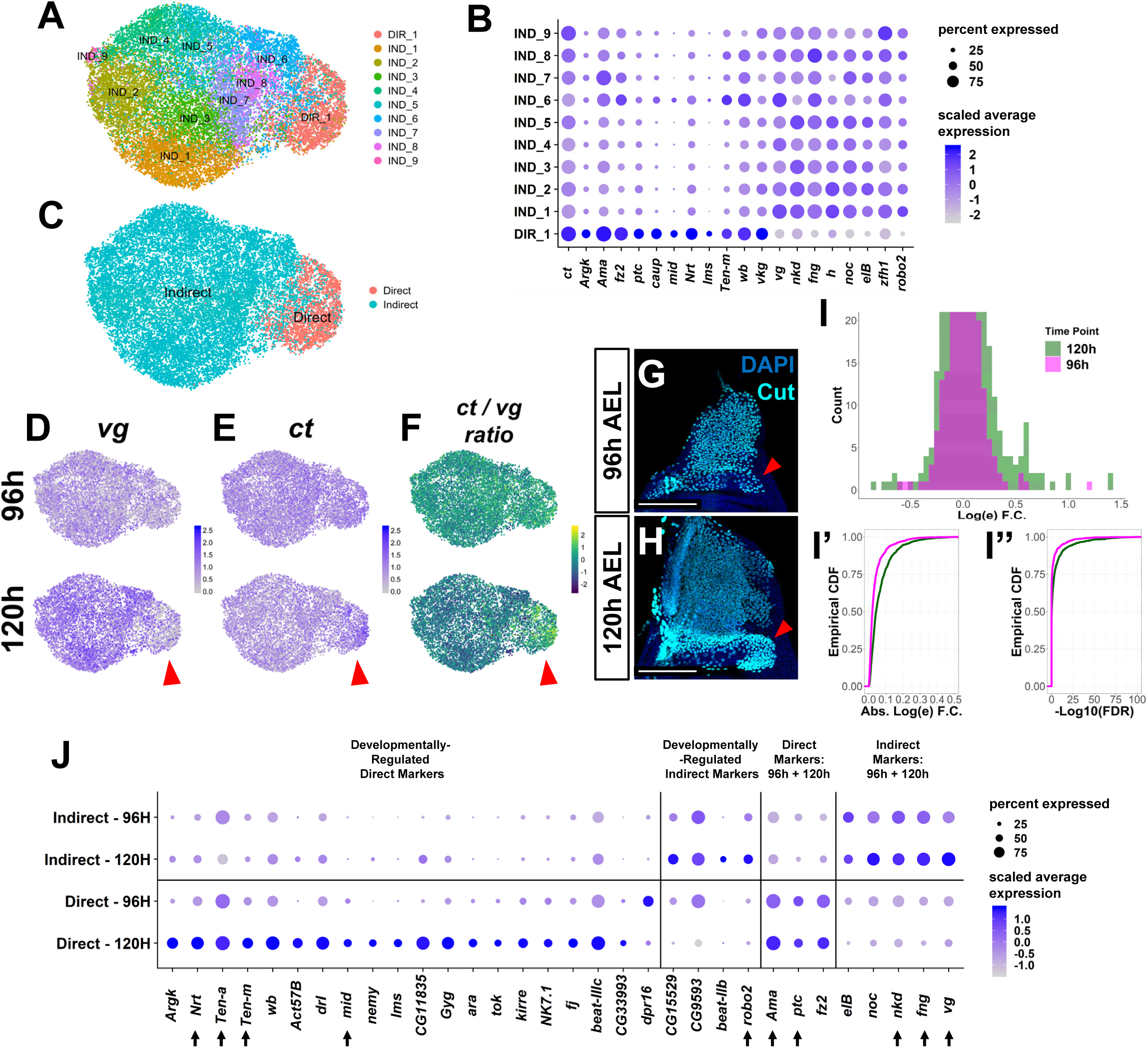
Cell atlas of the developing AMPs. (**A**) UMAP of AMPs, with cells colored by Seurat cluster identities after cell cycle and cell sex stratification has been corrected as detailed in **Figure 3 - figure supplements 1 and 2**. (**B**) Dot plot of AMP clusters showing markers of direct and indirect AFM precursors. The canonical marker genes *ct* and *vg* are shown, in addition to other genes that displayed differential expression between cluster DIR_1 and all other clusters (FDR < 0.05, see **Materials and Methods**). Note the higher expression of *ct* and lower expression of *vg* in DIR_1 compared to other clusters. When performing differential expression analysis on DIR_1 vs. all other clusters, there was a natural-log fold-change of 0.28 and −0.49 for *ct* and *vg*, respectively (FDR < 0.05 for both genes; positive natural-log fold-changes indicate higher expression in direct AMPs, negative values indicate higher expression in indirect AMPs) (**C**) UMAP of AMP single-cell data, as in (**A**), with cells colored by classification into putative precursors of direct or indirect AFMs. Clusters were assigned “direct” and “indirect” identity by relative expression levels of the marker genes *vg* and *ct*, as shown in **B**. (**D-F**) UMAPs separated by developmental time points showing the expression of canonical markers of indirect and direct AMPs, *vg* (**D**) and *ct* (**E**), with panel **F** showing a UMAP with each cell colored to indicate the ratio of *ct* to *vg* expression. Red arrowheads highlight differential expression of these genes in the direct AMPs at 120h. At 96h, the natural-log fold-change between direct and indirect AMPs for *ct* and *vg* is 0.12 and −0.28, respectively (FDR < 0.05 for both). At 120h, the natural-log fold-change for *ct* and *vg* increases (in terms of magnitude) to 0.44 and −0.69, respectively (FDR < 0.05 for both) (**G-H**) Wing discs from 96h (**G**) and 120h (**H**) AEL, stained with anti-Cut. Red arrowheads indicate location of direct AMPs, identifiable at 120h by higher anti-Cut staining and inferred by location at 96h. Images are max projections across image slices containing AMPs. Note that elevated anti-Cut staining is observed in the direct AMPs only at 120h, which matches the UMAP (**E**). (**I**) Distributions of log fold-changes and FDR values between direct and indirect AMPs at 96h (magenta) and 120h (green), characterizing the transcriptome-wide differences between cell types. The distributions only include variable genes (as determined by Seurat) that show consistent differential expression (i.e., same direction of fold-change) between batches. Natural-log fold-changes are calculated from the average fold-change across replicates, and the FDR values reported are the maximum (most conservative) FDR over all replicates (see **Materials and Methods**). Panel **I** displays the log fold-change distribution as a histogram, with the height of the y-axis limited to 20 for better visualization. For a given gene, positive magnitude log fold-changes indicate higher expression in the direct AMPs and negative magnitude log fold-changes indicate higher expression in the indirect AMPs, relative to each other. Panels **I’** and **I’’** display the log fold-change and FDR distributions, respectively, as empirical cumulative density functions (ECDF). These ECDF plots are calculated as the percentage of variable genes (y-axis) below a particular log-fold change magnitude threshold (**I’**) or FDR threshold (**I’’**). Note that at 120h (green) relative to 96h (magenta), the variable genes display more significant differential expression (lower FDR) and higher fold changes between direct and indirect AMPs. (**J**) Dot plot of marker genes of direct- or indirect-flight muscle precursors, with data separated by developmental time points to investigate temporal dynamics. Genes are grouped in the following manner: (1) genes that are differentially-expressed at higher levels within direct AMPs only at one time point (“Developmentally-Regulated Direct Markers”), (2) genes that are differentially-expressed at higher levels within indirect AMPs only at one time point (“Developmentally-Regulated Indirect Markers”), (3) genes that are differentially-expressed at higher levels within direct AMPs at both time points (“Direct Markers: 96h + 120h”), and (4) genes that are differentially-expressed at higher levels within indirect AMPs at both time points (“Indirect Markers: 96h + 120h”). Arrows highlight genes that are discussed in the main text. Color scales for UMAPs correspond to normalized (by total UMI) counts on a natural-log scale. Microscopy scale bars = 100 μm.

The AMPs are known to differentiate into either direct or indirect flight muscles of the adult fly (Bate, 1993; Roy & VijayRaghavan, 1999; Sudarsan et al., 2001). The precursors of these two populations can be identified by their location within the tissue, and are canonically classified by their relative expression of two transcription factors, Vestigial (Vg) and Cut (Ct), at the late third-instar larval (L3) stage (corresponding to our 120h time point) (Sudarsan et al., 2001). The precursor cells of the indirect flight muscle are localized more dorsally (closer to the wing disc stalk) and display relatively high Vg and low Ct protein expression (Sudarsan et al., 2001). The direct flight muscle cell precursors are localized more ventrally (closer to the wing hinge) and are identifiable by little or no Vg and high Ct protein expression (Sudarsan et al., 2001). We examined the expression levels of these two genes in our AMP cells, after they were grouped using unsupervised clustering (**Figure 3A, B**). We found that one cluster was characterized by high levels of *ct* and low levels of *vg* (**Figure 3B, D-F**). Conversely, all other clusters displayed relatively elevated levels of *vg* and low levels of *ct*. Based on this distinction, we classified clusters as representing the direct or indirect AMPs (**Figure 3C**), obtaining 17,604 indirect AMPs and 2,281 direct AMPs.

Classification of direct and indirect AMP cells based on *vg* and *ct* expression enabled us to identify additional genes that are differentially expressed between the two canonical AMP cell types (**Figure 3B** and **Materials and Methods**). These genes include *Zn finger homeodomain 1* (*zfh1*), *hairy* (*h*), and *naked cuticle* (*nkd*) for indirect AMPs and *Amalgam* (*Ama*) and *wing blister* (*wb*) for direct AMPs, which were also identified in a recent single-cell study on the AMPs (Zappia et al., 2019). Some of these identified genes were expressed at higher levels within certain subsets of indirect AMP, such as the transcription factors *h* and *rotund* (*rn*) that jointly mark a subset of indirect AMPs (**Figure 3B** and **Figure 3 - figure supplement 3**). Using a *rn-Gal4* line, we confirmed the expression of *rn* in a subset of AMPs nearest to the stalk of the wing disc (**Figure 3 - figure supplement 3**), which largely overlaps with the pattern identified for the gene *h* via antibody staining (Zappia et al., 2019). Potentially, *h* and *rn* may separate indirect AMPs into subsets that contribute to different muscle fibers, although such a role has not been described. More generally, these observations demonstrate that our AMP cell atlas can identify more refined partitions than the standard two groups based on *vg* and *ct* expression.

From mid to late 3rd instar, AMP number increases from approximately 630 to 2500 cells per wing disc (Gunage et al., 2014). However, the transcriptomic changes in this time interval have not previously been studied. We examined the expression of the canonical AMP markers *ct* and *vg* at single-cell resolution at 96h and 120h. Interestingly, both genes displayed more dramatic differential expression between direct and indirect AMPs at 120h as compared to 96h (**Figure 3D-F**). We confirmed this observation by examining Ct protein levels, which appear uniform within all AMPs at 96h as compared to the enrichment within the direct AMPs at 120h (**Figure 3G, H**). Together, this suggests that the direct and indirect cell identity is established over this developmental window as the differential expression of Ct is only observed at 120h. We broadened our analysis to a transcriptome-wide view and observed more significant differential expression between cell types (direct and indirect AMPs) at 120h as compared to 96h (**Figure 3I-I’’**). Thus, the direct and indirect AMPs are becoming more transcriptionally distinct over this last 24 hours of larval development. Examples of genes that are predominantly expressed in either direct or indirect AMPs are shown in **Figure 3J**. While there are many genes that are developmentally-regulated, note that there are genes that show consistent expression at both time points, possibly indicative of pathways that are necessary both for the initial establishment and subsequent maintenance of AMP cell types. This includes several predicted targets of Wg signaling, specifically *naked cuticle* (*nkd*) and *vg*, and the modulator of Notch signaling *fng* (**Figure 3 - figure supplement 3**).

From 96h to 120h, we noticed the activation and refinement of expression of many genes previously implicated in axon guidance (**Figure 3J**). These include the receptor *roundabout 2* (*robo2*), whose ligand *slit* (*sli*) (Kidd et al., 1999) was detected at low levels within the AMPs (**Figure 4H**), as well as synaptic partner-matching proteins *Tenascin major* (*Ten-m*) and *Tenascin accessory* (*Ten-a*) (Hong et al., 2012). In particular, we confirmed that Ten-m protein increases in expression from 96h to 120h by staining discs with anti-Ten-m antibody (**Figure 3 - figure supplement 3**). We also found that the axon guidance gene *Neurotactin* (*Nrt*) (Hortsch et al., 1990) marks the direct AMPs only at 120h. Interestingly, the reported ligand for Nrt, encoded by the gene *Amalgam* (*Ama*) (Frémion et al., 2000), is expressed in primarily the direct AMPs at both 96h and 120h. These observations suggest that pathways known to function in axon guidance could also function in myoblast fusion or muscle morphogenesis, and their expression may need to be restricted to either direct or indirect AMPs prior to pupal development.

**Figure 4.**
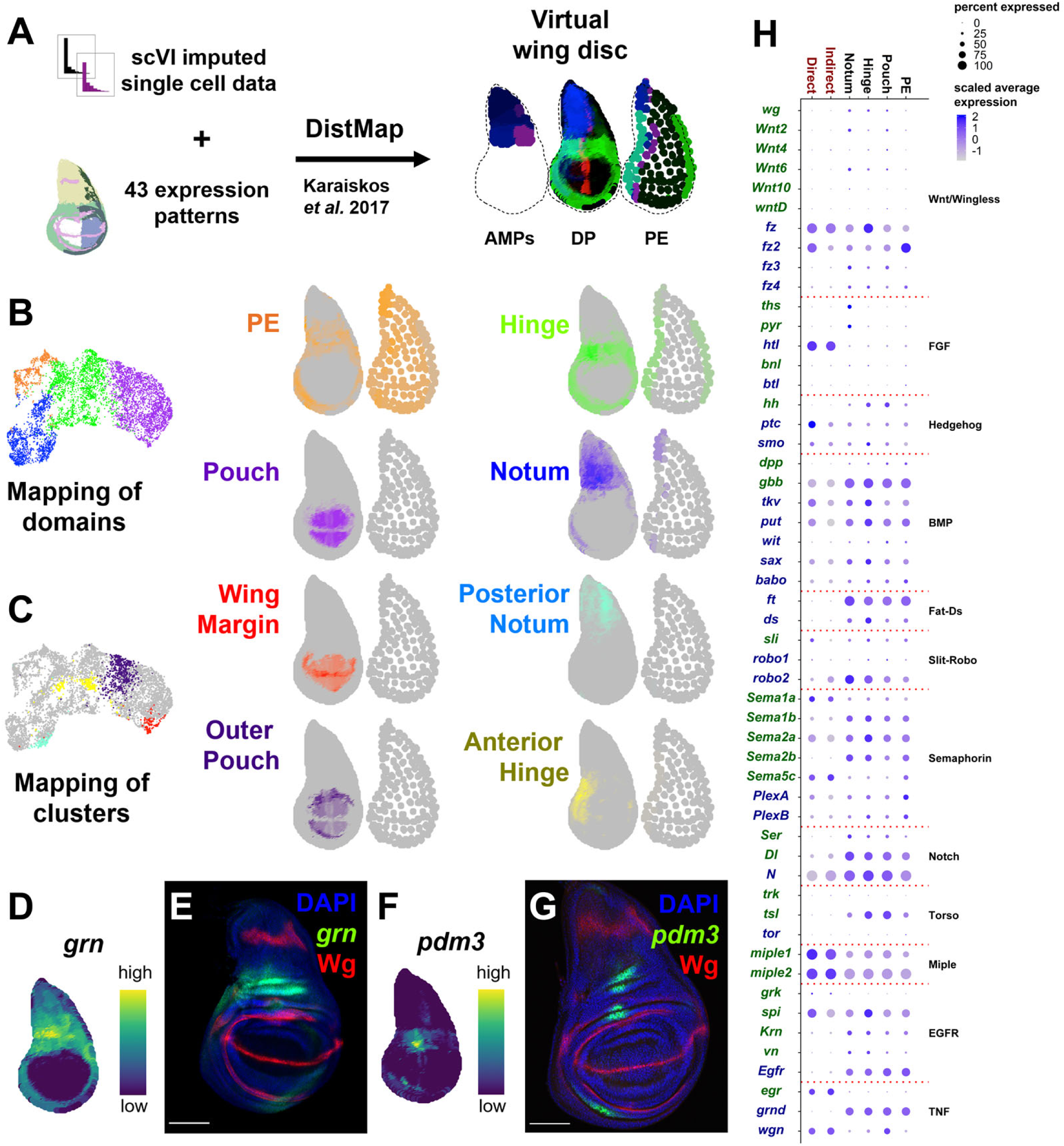
Virtual wing-imaginal disc and examination of receptor-ligand expression. (**A**) Schematic describing the creation of a three-layered virtual wing disc (AMPs, disc proper (DP), and the peripodial epithelium (PE)). In contrast to the DP which is mostly composed of columnar cells, much of the PE is composed of squamous cells with flattened nuclei, and it is therefore represented as an outline which contains large dots. We used scVI imputed gene expression values together with 43 reference gene expression patterns (see **Materials and Methods**) to statistically map our cells to locations within the reference geometry using Distmap (Karaiskos *et al.,* 2017). The virtual wing disc can be used to predict gene expression patterns (or virtual *in situ*), as shown for three example genes (*ptc* in red, *zfh2* in green, and *eyg* in blue). (**B, C**) Mapping of cells from the disc epithelium to the DP (left) and PE (right) layers of the virtual wing disc. (**B**) The cells from the epithelial domains mapped to the disc model. Stronger colors (orange, green, purple, and blue for PE, hinge, pouch, and notum cells, respectively) indicate higher predicted mapping; gray indicates low predicted mapping for cells. The UMAP is the same as in **Figure 2I**. (**C**) The cells from particular sub-regions are shown both on the UMAP and on the virtual wing disc; see **Figure 4 - figure supplement 1** for additional mapping of cell clusters. Stronger predicted mapping for cells in each cluster is indicated by higher levels of the corresponding color in the UMAP; gray indicates low predicted mapping. (**D, F**) Virtual *in situ* hybridizations to the DP layer as predicted by our virtual disc model where relative gene expression levels are shown on a scale from low (dark purple) to high (yellow). Predicted gene expression pattern of *grn* (**D**) and *pdm3* (**F**) in the epithelium disc proper. Neither of these two genes were used in building the disc model. (**E, G**) Late 3^rd^ instar wing-imaginal disc with transcriptional reporters for the genes *grn* (**E**) and *pdm3* (**G**). Note the relative similarity between the predicted expressions and transcriptional reporters. (**H**) Dot plot of expression of genes encoding receptors and ligands from pathways that were differentially-expressed in at least one cell type. X-axis: Cell groups. Disc epithelium cell types are colored black, AMP cell types are colored red. Y-axis: Genes are colored either blue or green depending on their annotation as encoding for a receptor or ligand, respectively. Color scales are on a natural-log scale. Microscopy scale bars = 100 μm.

### A three-layered virtual wing disc and expression of ligand-receptor pairs predict cell-cell interactions

In order to identify interactions between the epithelium and the AMPs, we took two approaches. First, to predict and visualize the location of gene expression within the wing disc, we generated a three-layered model, with gene expression levels inferred at different spatial positions along the AMP, disc proper, and peripodial epithelium layers. This enabled us to discover the expression of genes in regions of the disc epithelium that are closest to the AMPs. Second, we examined the expression of ligand-receptor pairs to identify those with complementary expression patterns between the disc epithelium and the AMPs.

To generate a three-layered transcriptomic map of the wing disc, we used the R package DistMap (Karaiskos et al., 2017) along with a manually curated set of published images of gene expression patterns (**Figure 4A** and **Materials and Methods**). The mapping of cells into their spatial origin was largely consistent with our marker-gene based cluster annotation (**Figures 2G** and **Figure 4 - figure supplement 1**). This included the four broad epithelial domains (notum, hinge, pouch, peripodial epithelium; **Figure 4B**) and more refined cell groups, such as the wing margin, posterior notum, outer pouch, and anterior hinge (**Figure 4C**). This virtual wing disc can be used to predict gene expression patterns by generating a “virtual *in situ”* (Karaiskos et al., 2017) (see **Materials and Methods**). To test how well our virtual wing disc would predict novel gene expression patterns, we predicted the expression of the genes *grain* (*grn*) and *pou domain motif 3* (*pdm3*), neither of which were included in the reference gene expression patterns used to generate the disc model. The virtual *in situs* predicted highest expression for *grn* in the anterior-dorsal region of the hinge (**Figure 4D**), and *pdm3* in small patches within the dorsal and ventral hinge regions (**Figure 4F**). These predicted patterns largely matched the expressions of transcriptional reporters for *grn* and *pdm3* within wing discs (**Figure 4E, G**), indicating that our virtual wing disc can successfully predict novel gene expression patterns. Thus, with the combination of our virtual wing disc and temporal cell atlas, we can localize gene expression and determine if their expression changes towards the end of larval development.

To look for potential cell communication between the different cell layers of the wing disc, we examined the expression of ligand-receptor pairs within the major domains of the disc epithelium and AMPs (**Figure 4H**) (for details on receptor-ligand pairs examined, see **Materials and Methods**). Consistent with previous work, we observed the expression of the ligand *wg* in the notum, as well as the genes encoding three other Wnt-family ligands *Wnt2*, *Wnt6*, and *Wnt10*. In contrast, *Wnt4* expression was mainly in the wing pouch (**Figure 4 - figure supplement 2**). Interestingly, we observed high levels of expression of the genes encoding for two FGF-family ligands, *thisbe* (*ths*) and *pyramus* (*pyr*) (Stathopoulos et al., 2004), in the notum region of the epithelium, whereas the gene encoding their receptor, *heartless* (*htl*) (Beiman et al., 1996), was specifically expressed in the AMPs. Similarly, the ligand *hedgehog* (*hh*) appears to be expressed only in the disc epithelium, while its receptor *patched* (*ptc*) and signal transducer *smoothened* (*smo*) are both expressed in the AMPs. Our detailed investigations of the FGF and Hedgehog pathways are presented in this study.

Additionally, our data also suggests that a signal could be transmitted from the AMPs to the epithelium via the ligand Eiger (Egr), which we find is transcribed in most AMPs (**Figure 4 - figure supplement 2**). Transcripts for its receptor Grindelwald (Grnd) were expressed in the epithelium. Likewise, several plexin and semaphorin genes (Tran et al., 2007) are expressed both in the epithelium and AMPs, and could be involved in bidirectional signals between the two cell types.

### FGF signaling from the epithelium creates a niche that regulates AMP number and localization

From our analysis of expression of ligand-receptor pairs, we identified two notum-specific ligands, *ths* and *pyr*, as being potential candidates involved in epithelium to AMP signaling (**Figure 4H**). Together, Ths and Pyr comprise the ligands for one of the *Drosophila* fibroblast growth factor (FGF) signaling pathways, and both interact with the receptor Htl (Stathopoulos et al., 2004) (**Figure 5A**). While *htl* is detected in nearly all of the AMPs, the three-layered disc map predicts that *ths* and *pyr* are selectively expressed in different regions along the notum of the disc epithelium (**Figure 5B-D**). Specifically, *ths* is localized to the most proximal region of the notum, while *pyr* has a broader expression pattern into the posterior hinge (compare **Figures 5C and 5D**). We used Gal4 lines to confirm the expression patterns of both *ths* and *htl* within the wing disc; *ths* was exclusively expressed in the proximal epithelial notum and *htl* was expressed in the AMPs (**Figure 5E, H**).

**Figure 5.**
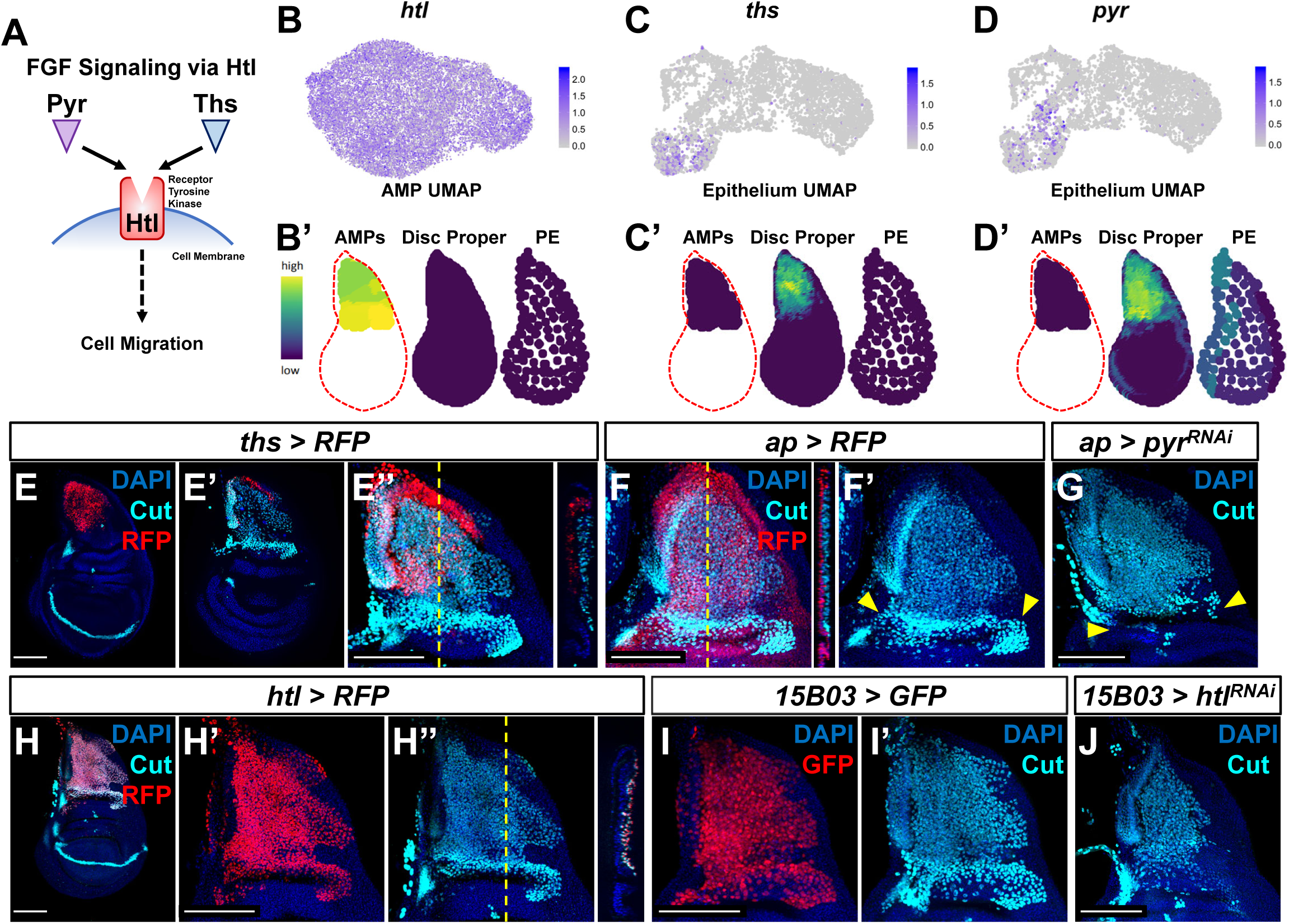
FGF knockdown reduces AMP number. (**A**) Cartoon of possible interaction between Pyr, Ths, and Htl. (**B-D**) Expression of *htl* (**B**), *ths* (**C**), and *pyr* (**D**) in the single-cell data. **B**, **C**, and **D** show UMAPs of these genes, in either the AMPs (for *htl*) or disc epithelium (for *ths* and *pyr*). **B’**, **C’**, and **D’** show virtual disc map predictions for the expression of these genes in the AMPs, disc proper, and peripodial epithelium. (**E-E’’**) Wing disc with *RFP* expression driven by *ths-Gal4* transgene. **E** is a max projection over image slices of the disc epithelium; **E’** is a max projection over image slices of the AMPs (visualized in cyan with anti-Cut staining). Orthogonal section (apical is left, basal is right) corresponds to the dashed yellow line in **E’’**. Note *ths-Gal4* drives expression specifically in the notum epithelium, but not in the AMPs. (**F, G**) Close up of the notum region from wing discs with *ap-Gal4* driving expression of *>RFP* (**F, F’**) *and >pyr^RNAi^* (**G**). Orthogonal section corresponds to dashed yellow line in **F**. Yellow arrowheads indicate expected location of direct AMPs, which can be identified in **F** by higher staining of anti-Cut. Note the loss of ventral- and posterior-localized AMPs following *pyr^RNAi^* in (**G**). (**H-H’’**) Wing disc with *RFP* expression driven by *htl-Gal4* transgene. **H** is a max projection image over all image slices. Orthogonal section corresponds to the dashed yellow line in **H’’**. Note *htl-Gal4* is expressed in the AMPs which express Cut. (**I, I’, J**) Close up of the notum region from wing discs with *15B03-Gal4* driving expression of *>GFP* (**I**) and *>htl^RNAi^* (**J**). AMPs visualized with anti-Cut. Note the reduction of AMPs, especially of the direct AMPs. Color scales for UMAPs correspond to normalized (by total UMI) counts on a natural-log scale. All notum images are max projections across image slices containing AMPs. Microscopy scale bars = 100 μm.

The genes *htl*, *ths*, and *pyr* have been studied extensively in the context of *Drosophila* embryogenesis. There, they are known to influence mesoderm spreading along the embryonic ectoderm and formation of cardiac progenitor cells (Beiman et al., 1996; Stathopoulos et al., 2004; Kadam et al., 2009). Notably, in *htl* mutants and in *ths* and *pyr* double mutants, mesoderm cells are still present within the embryo, but they accumulate in multilayered arrangements instead of a monolayer along the ectoderm (as observed in wild-type embryos) (Beiman et al., 1996; Stathopoulos et al., 2004; Kadam et al., 2009). These observations suggest that FGF signaling is primarily needed for proper mesoderm spreading, and possibly not for cell proliferation and survival. By analogy, Ths and Pyr may regulate the localization of AMPs in underlying the epithelial notum.

To examine the consequences of interfering with FGF signaling within the larval wing disc, we perturbed the expression of Pyr and Htl. To disrupt Pyr expression within the epithelial tissue, we utilized an *apterous* (*ap*) Gal4 driver (*ap-Gal4*) that expresses in the entire dorsal compartment of the disc proper, including all of the epithelial cells that overlie the AMPs (**Figure 5F**). Expressing an RNAi that targets *pyr* with this *ap-Gal4* driver resulted in a reduction in AMPs, primarily observed in the more ventral- and posterior-localized AMPs (**Figure 5G, compare with 5F**). A likely explanation for this result is that *pyr* knockdown within the wing disc restricts AMP survival and/or proliferation to the Ths-expressing region of the dorsal notum, as Pyr and Ths have been noted to have partially redundant functions (Stathopoulos et al., 2004; Kadam et al., 2009). We next tested if the Ths and Pyr receptor Htl is required within the AMPs. Using the AMP-specific driver *15B03-Gal4* (**Figure 5I**), we expressed an RNAi for *htl* and observed an obvious decrease in the number of AMPs following knockdown of FGF signal transduction (**Figure 5J**). Altogether, we conclude that FGF signaling between AMPs and the disc epithelium is necessary for proper AMP survival and/or proliferation.

To further determine if the location and level of FGF signaling controls the position and number of the AMPs, we assessed the effects of overexpressing the FGF ligands. First, we used a *dpp-Gal4* driver that is expressed in a stripe of cells just anterior to the anterior-posterior compartment boundary of the epithelium, including in the notum (**Figure 6A**). Expression of *pyr* in this domain caused a massive increase in the number of AMPs, not just adjacent to the epithelial notum (the AMP domain under wild-type conditions), but throughout the entire *dpp* domain (**Figure 6B**). Specifically, AMPs were observed along the entire stripe of ectopic *pyr* expression, all the way to the ventral hinge and even extending on the ventral side to the peripodial epithelium. From this, we conclude that FGF signaling can induce not just AMP proliferation, but also AMP spreading beyond the epithelial notum. These phenotypes were replicable when expressing the other FGF ligand, *ths*, with *dpp-Gal4* (**Figure 6C**). We further investigated the role of FGF signaling in AMP migration by generating a separate patch of *pyr* expression, discontinuous from the domain of endogenous expression. To this end, we ectopically expressed *pyr* within the wing pouch using the TRiP-Overexpression VPR toolkit (Lin et al., 2015) with a *nub* driver. We observed a large number of AMPs basal to the epithelium of the wing pouch (**Figure 6D, E**), suggesting that ectopic Pyr expression can recruit AMPs from a distance.

**Figure 6.**
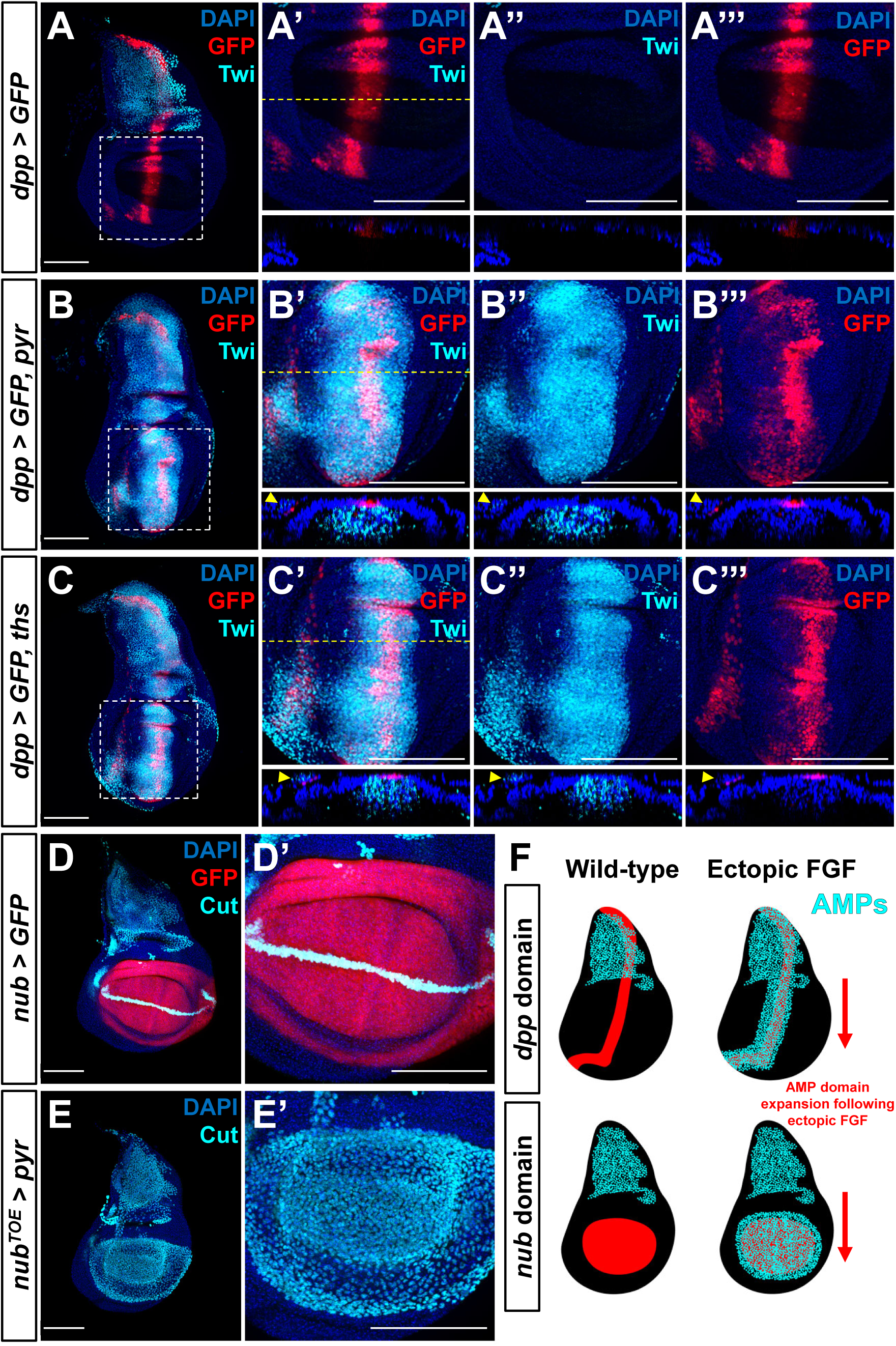
Ectopic FGF expression increases AMP number and migration. (**A-C**) Wing discs with *dpp-Gal4* driving the expression of *>GFP* alone (**A-A’’’**), or *>GFP* together with either *>ths* (**B-B’’’**) or *>pyr* (**C-C’’’**). The pouch region of each disc (corresponding to the dashed white boxes in **A**, **B**, and **C**) are shown at higher magnification in subsequent panels in each row. Orthogonal sections (apical is top, basal is bottom) correspond to dashed yellow lines in **A’, B’,** and **C’**. These discs (**A-C**) are stained with anti-Twi to visualize the AMPs. Yellow arrowheads indicate *dpp-Gal4* expression in the PE, which recruits AMP expansion to the PE surface when expressing either *>ths* or *>pyr*. (**D-E**) Wing discs with *nub-Gal4* driving the expression of *>GFP* (**D**) or *>dCas9^VPR^* (**E**), the latter of which can be used in conjunction with a guideRNA targeting an upstream sequence of the *pyr* transcriptional start site (*pyr^TOE.GS00085^*) to drive the overexpression of *pyr* in the wing pouch. These discs are stained with anti-Cut to visualize AMPs. Note that Cut is also expressed in the wing margin of the disc epithelium margin (seen as a band through the wing pouch in **D**). (**F**) Cartoon model for the effects of FGF overexpression on AMP growth. Ectopic expression of FGF ligands induces expansion of AMPs in a domain that broadly matches the pattern of FGF ligand expression. All wing disc images are max projections across all image slices. Microscopy scale bars = 100 μm.

These results regarding Htl, Ths, and Pyr point to two primary conclusions. First, increasing the levels of FGF signaling results in an increase of AMP numbers. This suggests a role of FGF signaling in providing trophic support for AMP survival and/or proliferation. Second, during normal development, the notum-specific expression of the ligands Ths and Pyr defines AMP localization. During wild-type development, this relationship between FGF signaling and AMP number could provide a mechanism by which the number and positioning of AMPs is matched to the size of the notum. Collectively, our experiments show that FGF signaling can both limit and induce the spread of AMPs along the disc epithelium, and as a result, FGF signaling effectively defines the AMP niche.

### Hedgehog signaling regulates gene expression in a subset of posterior localized AMPs

Our analysis of the expression of genes encoding ligand-receptor pairs also pointed to a possible role for Hh signaling (Lee et al., 2016) in a subset of the AMPs. We observed that *ptc*, which encodes the transmembrane receptor for the ligand Hh, is expressed at low levels in most AMPs and at a much higher level in a subset of the direct AMPs, while Hh signaling pathway components *smo* and *ci* are expressed in most AMPs at uniform levels (**Figure 7A-C** and **Figure 4H**). In the wing disc epithelium, cells within the posterior compartment are known to produce the ligand Hh, whereas cells of the anterior compartment are capable of receiving the Hh signal and activating a Hh-responsive signaling pathway (Jiang & Hui, 2008). The selective expression of Ptc in a subset of AMP therefore raises the hypothesis for transduction of the Hh developmental signal from the posterior compartment of the epithelium to a specific subset of myoblasts. However, while Hh signaling has been studied extensively in the context of the disc epithelium (reviewed by Lee et al., 2016), a function for the pathway in AMP development has not been described. Moreover, since we could not find evidence for expression in AMPs of *dpp,* the canonical target of Hh signaling in the disc epithelium, this raises the possibility that Hh signaling activates a different set of target genes in the AMPs.

**Figure 7.**
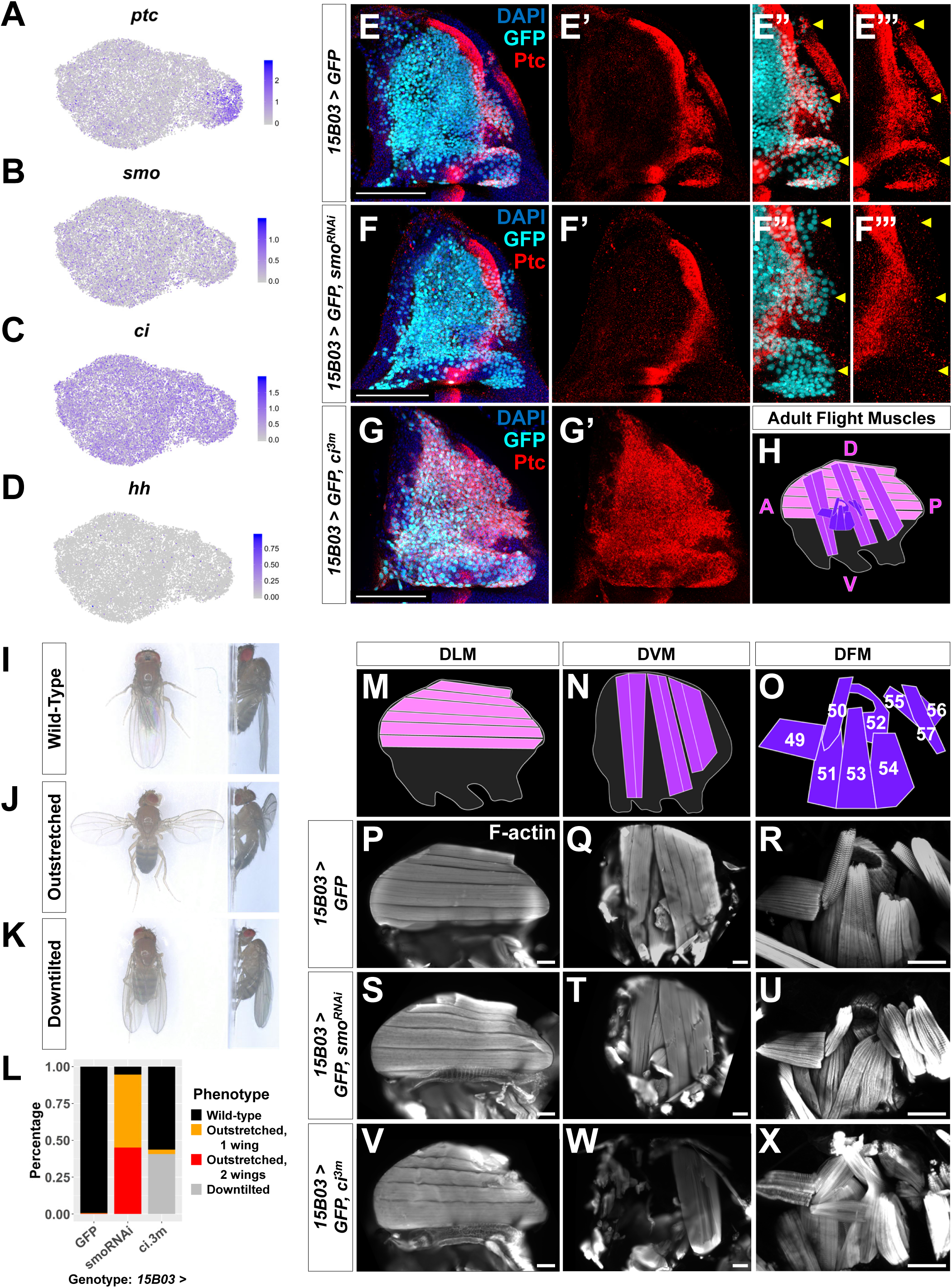
Hh signal from the disc patterns a subset of AMPs. (**A-D**) UMAPs of *ptc* (**A**), *hh* (**B**), *smo* (**C**), and *ci* (**D**) expression in AMPs. (**E-G**) Close up of the notum region of wing discs stained for anti-Ptc (red) with AMP-specific *15B03-Gal4* driving *>GFP* alone (**E**), or *>GFP* together with either *>smo^RNAi^* (**F**) or *>ci^3m^* (**G**). In the control, note that the AMPs expressing Ptc are to the right of the epithelium Ptc stripe. Panels **E’’**, **E’’’**, **F’’**, and **F’’’** focus on posterior-localized AMPs. Yellow arrowheads indicate groups of posterior-localized AMPs with high anti-Ptc staining in the control, which are absent following knockdown of *smo*. Note that following the overexpression of *>ci^3m^* that Ptc is expressed in all of the AMPs (**G’**). (**H**) Schematic of adult flight muscle fibers within the thorax where the different muscle subtypes are differentially shaded. The DLMs are in light purple, the DVMs and in intermediate purple, and the DFMs are in dark purple. (**I-K**) Wing posture phenotypes observed in Hh-signaling perturbation experiments. Adult flies were imaged live and not anesthetized. (**I**) Wildtype posture, with wing blades folded along their dorsum. (**J**) Outstretched wing posture, where either one or both wings were always held perpendicular to the body axis. (**K**) Downtilted wing posture, with adults that hold their wings farther apart along their dorsum and tilted laterally downward. (**L**) Quantification of observed wing posture phenotypes under conditions of Hh-signaling perturbation within AMPs. All *UAS* lines were driven via *15B03-Gal4*. The number of adult flies assayed: *>GFP* = 341, >*smo^RNAi^* (Bloom. #43134) = 283, and *>ci^3m^* = 366. (**M-O**) Separate schematics of expected DLMs (**M**), DVMs (**N**), and DFMs (**O**) morphology. Numbers on DFMs represent the canonical labels for the different fibers. (**P-X**) Adult flight muscles (visualized with F-actin staining) from animals with *15B03-Gal4* driving *>GFP* alone (**P-R**), or *>GFP* together with either *>smo^RNAi^* (**S-U**) or *>ci^3m^* (**V-X**). DLMs are shown in **P**, **S**, and **V**; DVMs are shown in **Q**, **T**, and **W**; DFMs are shown in **R**, **U**, and **X**. Adult flight muscles in *>GFP* flies had similar morphology in all flies examined (23 DLMs, 15 DVMs, and 12 DFMs examined) (**R**). Adult flight muscles in *>smo^RNAi^* (Bloom. #43134) animals displayed abnormal DFMs in all 11 flies examined (11/11 had muscles 53 and 54 misaligned, and 7/11 had muscles 55, 56, and 57 malformed) (**U**), while DLMs and DVMs had relatively normal morphology (observable in 21/22 DLMs and 10/10 DVMs) (**S, T**). These *smo^RNAi^* results were replicable with multiple RNAi lines. Adult flight muscles in *>ci^3m^* animals had normal morphology in all 7 DLMs examined (**V**), whereas all 7 DVMs examined were either missing or severely disconnected (**W**) and all 4 DFMs examined appeared abnormal (**X**). Color scales are on a natural-log scale. All notum images are max projections across image slices containing AMPs. Microscopy scale bars = 100 μm.

To confirm that the Hh ligand is likely originating from the epithelium, we looked at *hh* expression within our cell atlas. We found that only 1.1% of AMPs expressed *hh* transcripts and that these cells were sparsely scattered through the different AMP cell subsets (**Figure 7D**), indicating that there is not a specialized *hh*-producing AMP subtype. Furthermore, we did not detect expression of *hh-Gal4* within the AMPs (**Figure 7 - figure supplement 1**), providing additional evidence that the AMPs are not the source of Hh ligand. In contrast, *hh* transcripts were detected in approximately 32% of cells in the epithelium, roughly the size of the Hh-producing posterior compartment (**Figure 7 - figure supplement 1** and **Figure 4H**). Thus, these observations support the hypothesis that Hh originates from the disc epithelium and suggests that the *ptc-*expressing AMP cells would be located close to the epithelial source.

We determined the location of the Ptc-expressing AMP cells with immunohistochemistry. We detected Ptc protein in a subpopulation of AMPs, localized primarily beneath the posterior compartment of the disc epithelium (**Figure 7E**). Consistent with our scRNAseq data, Ptc was observed mostly in a group of direct AMPs, but also in two smaller clusters that are located more dorsally among indirect AMPs. The position of these Ptc-expressing cells within the tissue suggests that Hh ligand secreted from the epithelial cells of the posterior compartment is regulating gene expression in a subset of AMPs.

To determine if Hh signaling is important for specifying a subpopulation of AMPs, we manipulated the Hh-signaling pathway specifically in the AMPs. We observed that the knockdown of *smo*, which would reduce Hh signaling, resulted in the loss of Ptc expression in AMPs (**Figure 7F**). This indicated that, as in the disc epithelium, Hh signal transduction within the AMPs is required to establish high Ptc expression in the posterior-localized AMPs. Conversely, the expression of an activated form of the transcription factor Ci (Ci^3m^), which is resistant to proteolytic cleavage (Price & Kalderon, 1999), induced high levels of Ptc protein expression in all AMPs (**Figure 7G**). Thus, all AMPs appear capable of responding to Hh, but during normal development, only AMPs with close proximity to the posterior compartment of the disc epithelium receive the signal. Of note, neither *smo^RNAi^*-knockdown nor *ci^3m^* overexpression caused obvious changes in the overall number of AMPs (**compare Figure 7E with Figures 7F, G**). This suggests that Hh signaling in the AMPs does not have a large effect on AMP proliferation or survival during larval development, but could potentially be important for cell-fate specification.

To investigate a possible role of Hh signaling in AMP cell fate specification, we examined adult flies after genetic perturbations. During the pupal phase, the AMPs give rise to three distinct muscle fiber types within the adult thorax: dorsal longitudinal muscles (DLM), dorsoventral muscles (DVM), and direct flight muscles (DFM) (Figure 7H). While DLMs and DVMs are indirect flight muscles that generate the mechanical movement required for flight by compressing the thorax, the DFMs are responsible for flight steering by fine-tuning the position of the wing blades (reviewed by Bate, 1993). Both DLMs and DVMs are formed from indirect AMPs, whereas the direct AMPs develop into the DFMs. Control adults displayed wild-type posture (**Figure 7I**), while after Hh signaling was downregulated in AMPs with *smo^RNAi^*, we found that a majority of adults displayed an “outstretched” wing posture phenotype (**Figure 7J, L**). When *ci^3m^* was expressed in all AMPs, we observed that many adults displayed a “downtilted” wing posture (**Figure 7K, L**). These wing posture phenotypes were reproducible with multiple *smo^RNAi^* and in both sexes (**Figure 7 - figure supplement 2**). Adults with either the outstretched or downtilted phenotypes were incapable of flight. These observations suggest a vital role of Hedgehog signaling within the AMPs for the formation of functional adult flight muscles.

To examine if Hh signaling perturbations affected the structure of adult muscle fibers, we dissected adult thoraxes. Proper morphology of all three types of adult muscle fibers was observed in our controls (**Figure 7M-R**). However, when we reduced *smo* expression in the AMPs, we observed a misalignment of the DFM fibers (**Figure 7U**). In particular, the more posterior DFMs 52-57 (Miller, 1950; Bate, 1993; Ghazi et al., 2000) displayed improper position and overall disorganization (**Figure 7 - figure supplement 2**). Muscle 53, for example, inappropriately projects to the dorsal attachment site of muscle 54. In contrast, the DLM and DVM muscle fibers appeared relatively normal (**Figure 7S, T**). In contrast, expression of *ci^3m^* within the AMPs caused severe defects in the indirect flight muscles; most of the DVMs were eliminated (**Figure 7W**) and the DFMs were often severely disorganized and malformed (**Figure 7X**). Additionally, DFM muscle 52 was often missing or malformed. Importantly, muscle 51, which is known to arise from a separate group of AMPs not associated with the wing disc (Lawrence, 1982), is unaffected by these manipulations. Once again, the DLMs had no noticeable defects (**Figure 7V**). A possible explanation for this is that the DLMs unlike the other adult flight muscles do not arise in *de novo*, but rather by the fusion of AMPs with histolyzing larval muscles that act as templates (Fernandes et al., 1991). Overall, our data suggest that a normal level of Hh-signaling is important for proper specification of a subset of the direct AMPs and that excessive Ci activity can also perturb the development of some of the indirect flight muscles.

### Neurotactin and Midline are AMP-specific downstream targets of Hedgehog signaling

Since the canonical target of Hh signaling in the disc epithelium, *dpp*, is not expressed in the AMPs we looked for genes that correlated with *ptc* in our AMP cell atlas. Expression of both *Neurotactin* (*Nrt*) and *midline* (*mid*) displayed relatively high correlation with that of *ptc* (Pearson correlation of 0.34 and 0.44 for *ptc* / *Nrt* and *ptc* / *mid*, respectively). Mid, also known as Neuromancer 2, is a T-box transcription factor most related to mouse Tbx-20 (Buescher et al., 2004) that regulates cell fate in the developing nervous system (Leal et al., 2009). Nrt encodes a single-pass transmembrane protein expressed on the cell surface (Hortsch et al., 1990). Dimers of the secreted protein Amalgam (Ama), which are expressed in the direct AMPs, are able to bind to Nrt on two different cells and promote their adhesion (Frémion et al., 2000; Zeev-Ben-Mordehai et al., 2009).

We observed that both *Nrt* and *mid* expression increased dramatically in direct AMPs from 96h to 120h based on our single-cell data (**Figure 8A, B**) and antibody staining (**Figure 8D-G**), whereas *ptc* is more stably expressed (**Figure 8C, H, I**). Surprisingly, while Nrt and Mid expression patterns included all posterior-localized direct AMPs, the expression of both genes extended into anterior-localized direct AMPs which do not express Ptc.

**Figure 8.**
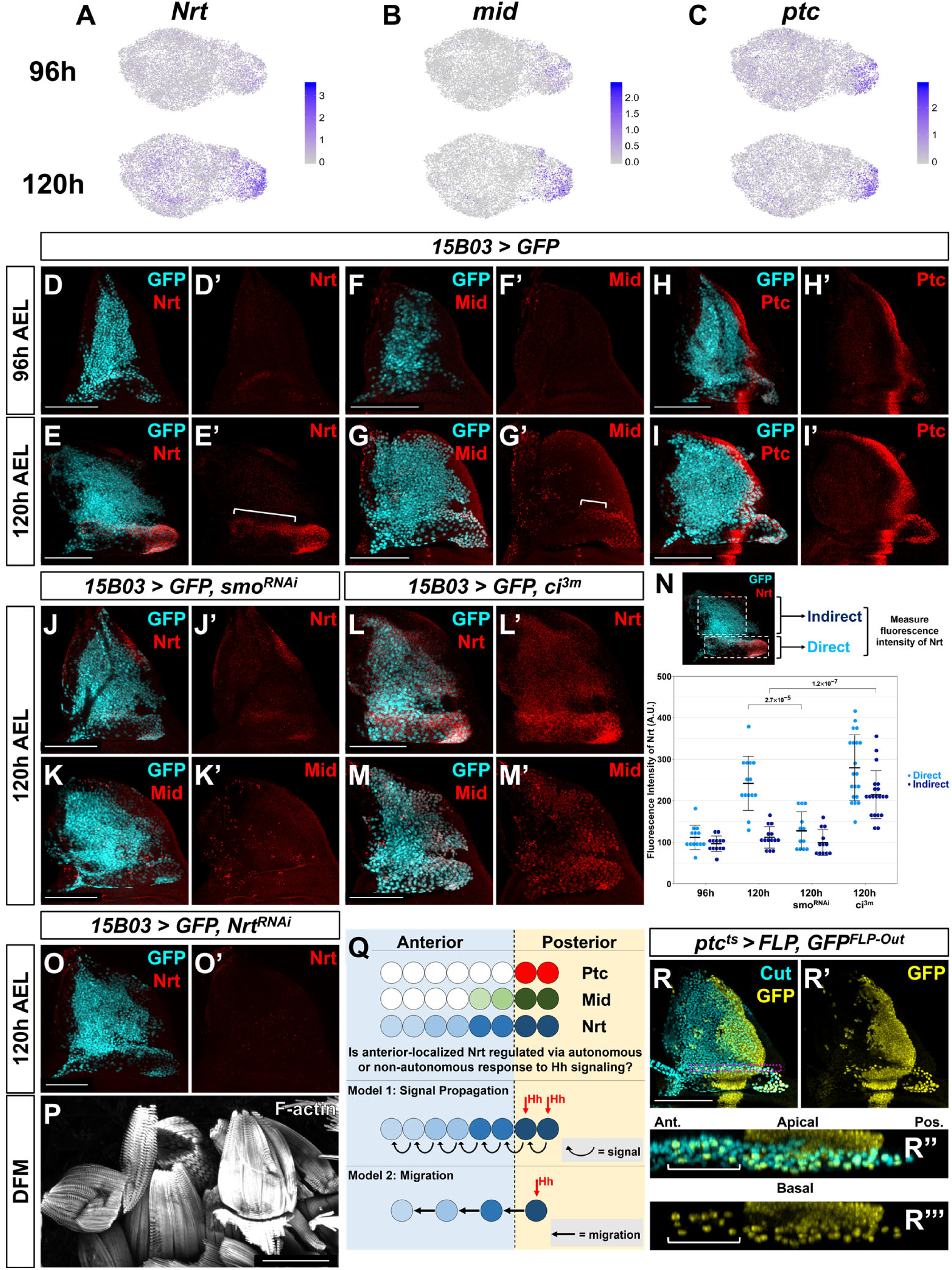
Nrt and Mid are downstream Hh-pathway targets in the AMPs. (**A-C**) UMAPs of *Nrt* (**A**), *mid* (**B**), and *ptc* (**C**) expression at 96h and 120h in AMPs. Note the increase in expression for *Nrt* and *mid* from 96h to 120h, whereas *ptc* expression is relatively unchanged. Within the direct AMPs, *Nrt*, *mid*, and *ptc* increased by a natural-log fold-change of 1.09, 0.47, and 0.12, respectively. (**D-I**) Wing discs stained with anti-Nrt at 96h (**D**) and 120h (**E**), anti-Mid at 96h (**F**) and 120h (**G**), and anti-Ptc at 96h (**H**) and at 120h (**I**). AMPs are visualized by expression of GFP (cyan) via *15B03-Gal4* driver. Note the negligible staining of anti-Nrt and anti-Mid in AMPs at 96h, mirroring the scRNAseq expression results. (**J-M**) Wing discs expressing either >*smo^RNAi^* (**J, K**) or >*ci^3m^* in AMPs via *15B03-Gal4* driver, along with GFP to visualize AMPs (cyan). Discs are either stained with anti-Nrt (**J, L**) or anti-Mid (**K, M**). Note that the knockdown of *smo* prevents the expression of Nrt and Mid in the direct AMPs and that the overexpression of *ci^3m^* leads to ectopic expression in the indirect AMPs. (**N**) Quantification of anti-Nrt staining within direct and indirect AMPs. Average fluorescent intensity was calculated within the boxed regions shown at the top. The graph shows binned values of average fluorescent intensity. P-values were calculated from unpaired t-tests and error bars indicate standard deviation. Note that the increased expression of Nrt specifically in the direct AMPs at 120h is not observed following *smo* knockdown. Number of discs examined: 96h wild-type: 13, 120h wild-type: 14, 120h *smo^RNAi^*: 12, 120h *ci^3m^*: 20. The overexpression of activated *ci* increases Nrt expression in both the direct and indirect AMPs. (**O**) Wing disc expressing *>Nrt^RNAi^* in AMPs via *15B03-Gal4* driver, stained with anti-Nrt at 120h. Note that knockdown of *Nrt* in the AMPs eliminates Nrt staining in the AMPs (**O’**). (**P**) DFMs in adults where *15B03-Gal4* drives *>Nrt^RNAi^*. Note the enlarged posterior DFMs, specifically muscles 55, 56, and 57 (similar phenotypes were observed in all 5 flies examined). (**Q**) Models explaining the protein expression of Hh-signaling targets in anterior-localized AMPs. In Model 1, posterior-localized AMPs receive Hh from the epithelium and propagate a secondary signal to anterior AMPs, inducing Nrt and Mid expression. This propagated signal is weaker in cells farther from the source of Hh. In Model 2, posterior-localized AMPs receive Hh from the epithelium and then migrate anteriorly (either a result of active cell movement or due displacement caused by proliferation). Anterior-localized AMPs quickly degrade Ptc protein, but Hh-signaling targets Nrt and Mid perdure longer. (**R**) Lineage tracing of AMPs that have previously expressed *ptc* in the larval stage. Expression of GFP in wing discs marked by *ptc-Gal4, tub-Gal80^ts^, >FLP, Ubi-FRT-stop-FRT-GFP* (*ptc^ts^>FLP, GFP^FLP-Out^*). Cells that expressed high levels of *ptc-Gal4* while at the non-permissive temperature for Gal80^ts^ (30 °C) will be permanently labeled by GFP expression. The temperature shift from 18 °C to 30 °C was done at 5 days AEL for 24h and the larvae were dissected at late 3rd instar. AMPs were visualized with anti-Cut staining. Orthogonal max projection is shown in **R’’** and **R’’’**, corresponding to the dashed purple box in **R**. White brackets in **R’’** and **R’’’** indicate anterior-localized AMPs. Note that a subset of the anterior labeled AMPs (bracketed) expresses GFP. Color scales are on a natural-log scale. All notum images are max projections across image slices containing AMPs. Microscopy scale bars = 100 μm.

Due to their high expression levels in the posterior-localized AMPs, we hypothesized that expression of Nrt and Mid, at least in the posterior AMPs, was influenced by Hh signaling. To determine if *Nrt* and *mid* are Hh signaling targets, we examined if perturbing the Hh pathway within the AMPs would alter their expression at 120h AEL. The knockdown of *smo* and consequent reduction in Hh signaling resulted in a dramatic decrease in both Nrt and Mid expression in the direct AMPs (**Figure 8J, K, N**). Remarkably, this was observed in both the posterior- and anterior-localized AMPs alike. In contrast, *smo* knockdown did not affect Ct expression in the direct AMPs (**Figure 8 - figure supplement 1**), which indicates that Hh signaling is not essential for direct AMP identity, as assessed by high Ct expression, but is key for additional cell fate specification. We tested if increased Hh signaling would be sufficient to induce expression of Nrt and Mid by driving *ci^3m^* in the AMPs. This resulted in the ectopic expression of both Nrt and Mid in all of the AMPs, although expression was higher within direct AMPs (**Figure 8L-N**). In contrast to the AMPs, we observe very low correlation between *ptc* and either *Nrt* or *mid* within the cell atlas of epithelium. These experiments suggest that Hh signaling is important for proper patterning of the AMPs and that Nrt and Mid are AMP-specific targets of the Hh pathway.

To evaluate the functional consequences of reducing Mid and Nrt expression, we used RNAi to knockdown their expression. The RNAi line directed against *mid* did not reduce Mid protein levels. However, we were able to reduce Nrt levels using RNAi (**Figure 8O**), which resulted in defects in the more posterior DFMs albeit not as severely as knockdown of *smo* (**Figure 8P**). This result shows that the Hh-signaling target Nrt is critical for proper DFM development.

We were, however, puzzled by the anterior expression of both Nrt and Mid, since these AMPs were outside the region of Hh signaling, at least as assessed by Ptc expression (**Figure 8Q**). We considered two models to explain this discrepancy. In the first model, Hh signaling in the posterior AMPs would induce the expression of a secondary signal that spreads to the anteriorly-localized AMPs to activate the expression of *Nrt* and *mid*. In the second model, the anterior-localized AMPs would have experienced Hh-signaling earlier in development by past proximity to the Hh-producing posterior compartment of the disc epithelium. The current location of these cells would be due to either an active anterior migration or a stereotyped pattern of cell displacement as a result of proliferation (**Figure 8Q**). To test whether anterior-localized AMPs had past activation of the Hh-signaling pathway (i.e., the second model), we used a lineage-tracing method to identify cells that had previously expressed *ptc*. *ptc-Gal4* was used to drive the expression of *UAS-FLP* which, in turn, elicited a recombination event that resulted in constitutive GFP expression. When this was done throughout all of development, we observed that all AMPs had expressed *ptc* at some point. To exclude embryonic expression from our analysis, we narrowed the window of labeling to the 3rd instar using a temperature-sensitive Gal80 to restrict the activity of *ptc-Gal4* (see **Materials and Methods**). We then observed that all the direct AMPs underlying the posterior compartment of the epithelium expressed GFP, as well as a trail of anterior AMPs that recapitulate the domain of anti-Nrt staining (**Figure 8R**). These results indicate that these anterior GFP-positive AMPs were descended from high *ptc-*expressing cells, and that past activation of the Hh-pathway is likely to be responsible for the expression of both anterior- and posterior-localized Mid and Nrt. These observations are consistent with cell migration as outlined in model 2, which highlights that past cell-cell interactions impact gene expression patterns later in development.

## Discussion

Organs are typically composed of many different types of cells, often including cells from different germ layers. Signals are exchanged between subsets of cells in ways that are restricted both spatially and temporally. With the eventual goal of obtaining as complete an understanding as possible of all of the signals exchanged between cells during the development of the *Drosophila* wing disc, we have generated cell atlases of the wing disc from two time points in larval development. We have catalogued temporal transcriptional changes that occur both globally and in subsets of cells. Additionally, we generated a way of visualizing gene expression simultaneously in the three layers of the wing disc. This has enabled us to capture the diversification of cell types that occurs in both the disc epithelium and the underlying AMPs over the same time interval and to identify and characterize ligand-receptor interactions that occur between different types of cells in spatially restricted domains.

### Heterogeneity and Diversification of Cell Types

Until recently, most genome-wide data on gene expression in the wing disc was derived from bulk RNA-seq or microarray experiments (Arbeitman et al., 2002), that used either entire discs or specific regions of the disc that had been physically fragmented. While such studies have contributed significantly to our understanding of temporal changes in gene expression, they did not provide the spatial resolution to parse expression changes that occur in different subpopulations of cells. The expression levels of genes in specific subsets of cells have previously been visualized using RNA *in-situ* hybridization experiments and reporter constructs such as MiMIC lines (Venken et al., 2011) and the FlyLight collection (Jory et al., 2012). However, the readouts of these experiments are difficult to quantify and do not provide an easy comparison of the relative level of expression of different genes. In contrast, the data obtained from scRNAseq experiments can provide both spatial and temporal information that give us a better understanding of how cells diversify and then stabilize their transcriptomes during development, and point to ways in which they interact with each other.

One remarkable observation from our data is that spatial positioning within the wing disc is highly informative of the transcriptional state of cells. In particular, the proximo-distal axis of the disc epithelium is one of the primary stratifying features within our single-cell data. Cell clusters, as identified by our analysis of the disc epithelium data, can be efficiently grouped together to define different sub-regions in the notum, hinge, pouch, and peripodial epithelium. In contrast, although the cells of the anterior and posterior compartments have been separated by lineage since early in embryogenesis, we observe less differential expression between the two compartments as compared to differential expression between proximodistal regions (notum, hinge, and pouch). Many clusters within the pouch, in particular, are composed of cells from both compartments. Thus, position along the proximodistal axis has a far greater influence on the transcriptome of a cell than its ancestry. This observation may also explain the challenge faced by studies that have aimed to find differences between the two compartments in order to explain why these two populations of cells remain segregated (Umetsu et al., 2014).

In the epithelium, we observe that most of the major cell types observed at late L3 (120h AEL) are already present at mid L3 (96h AEL). However, the transcriptomes of the two major populations of AMPs, those that give rise to the direct and indirect flight muscles, diverge significantly during this time interval. At 96h, AMPs appear to be in a relatively naive state; canonical markers for the direct and indirect flight precursors, *ct* and *vg*, both show relatively uniform expression at the mid L3 stage. At the late L3 stage, we observe more distinguishable differences between the transcriptomes of direct and indirect cell types. Both *ct* and *vg* have greater differential expression in the AMPs at this time point, and we identify the rise in expression of AMP-specific Hh-pathway targets *Nrt* and *mid*. Furthermore, the direct and indirect AMPs become marked by the differential transcription of a panel of axon-guidance genes. We speculate that these genes may facilitate the process of AMP migration to pupal fusion sites within the thorax, as well as aid in the fusion of direct and indirect AMPs with their corresponding cell types.

As the larva progresses through L3 and approaches the onset of metamorphosis, there is an increase in the circulating levels of the steroid hormone ecdysone. Our data show that ecdysone target genes can be differentially activated in different populations of cells. For example, the genes *ImpE2* and *Hr4* are upregulated in both epithelial cells and myoblast, while others such as *ImpE1* and *ImpE3* are mostly upregulated in epithelial cells alone. The mechanisms that modulate hormonal signaling within individual cell types represents an interesting area for future study.

### FGF signaling regulates the number and location of AMPs

We have shown that localized expression of the two FGF-family ligands Ths and Pyr in the notum restricts the AMPs to this region. Ths and Pyr from the epithelial cells activate the signaling pathway downstream of Htl in the underlying AMPs. Ths and Pyr have previously shown to regulate the spreading of mesodermal cells during embryogenesis but, to our knowledge, a role for these ligands in regulating myoblast numbers was not previously appreciated. We show that antagonizing this pathway reduces AMP numbers and that increased expression can induce a dramatic overproliferation of the AMPs. Thus, the level of Ths and Pyr secreted by epithelial cells in the notum could provide sufficient trophic support to generate the appropriate number of AMPs during normal development. While this work was in preparation, another group independently showed that the *ths-Gal4* line is expressed in the epithelium of the notum and that reducing *ths* function reduces AMP numbers (Vishal et al., 2020).

We have also demonstrated that ectopic expression of Ths or Pyr can draw AMPs out of the notum region, all the way to the ventral hinge and around the ventral edge of the disc proper onto the peripodial epithelium. Conceivably, expression of Ths and Pyr in the notum could be responsible for drawing AMPs into the disc epithelium at earlier stages of development. Indeed, even the expression of Pyr in the wing pouch, which is separated from the notum by the dorsal hinge, was sufficient to promote colonization of the pouch region by AMPs. This observation raises the possibility that these FGF proteins could act as long-range chemoattractants or that the myoblasts might have processes that could sense FGF proteins at considerable distances.

### Instructive Hedgehog signaling from the epithelium to the myoblasts

We have shown that the anteroposterior patterning of the disc epithelium is important for proper specification of gene expression within the underlying AMPs. Previous studies have shown that the stripe of Wg expression in the notum promotes the expression of Vg in the underlying myoblasts and that Notch signaling switches myoblasts from a symmetric to an asymmetric mode of cell division (Gunage et al., 2014). Because Hh has a short range of action, either due to its diffusive properties or because it is taken up by receiving cells by projections known as cytonemes (Parchure et al., 2018), only the AMPs underlying the posterior compartment respond to Hh by the upregulation of target genes. We show that all AMPs are potentially capable of transducing the Hh signal since they express the *hh* receptor ptc at low levels, the signal transducer *smo*, and the transcription factor *ci*.

We also describe two AMP-specific Hh targets, *mid* and *Nrt.* While we currently do not know whether mid and Nrt are direct targets of Ci, both *Nrt* and *mid* do have consensus Ci-binding sites within potential regulatory regions. Nrt is a single-pass transmembrane protein. Its extracellular ligand, Amalgam, has more widespread expression in the direct myoblasts and is expressed at comparable levels at 96h and 120h AEL. Two molecules of Amalgam can form homodimers and each is capable of binding to Nrt on a different cell (Frémion et al., 2000; Zeev-Ben-Mordehai et al., 2009). Thus, an effect of Hh-induced expression of Nrt in a subset of the direct AFMs might be to promote aggregation of Nrt-expressing cells at a later stage of development.

An unexpected observation was that AMPs beneath the anterior compartment express Nrt and, to a lesser extent, Mid. This expression is dependent upon Hh-signaling since knockdown of *smo* blocks gene expression. Since Hh signaling is restricted to the AMPs underlying the posterior compartment, as assessed by Ptc expression, the expression of Nrt and Mid in more anterior AMPs is not easily explained. Although we cannot completely exclude the possibility that a second signal from posterior AMPs activates *mid* and *Nrt* expression in these cells, our lineage-tracing experiments favor a model where a subset of the direct AMPs are generated posteriorly and move anteriorly during the course of development. Such movement could be due to a process of active migration in response to hitherto unknown external cues or to displacement as a result of oriented cell division. Understanding the mechanistic basis of AMP migration would represent an exciting avenue of future research.

### Identification of other ligand-receptor interactions in the wing disc

Our data also points to other possible signaling events between the disc epithelium and the AMPs. Both *dpp* and the related ligand *glass bottom boat* (*gbb*) are expressed predominantly in the disc epithelium. Genes encoding their receptors *tkv*, *put*, and *sax* are expressed in AMPs, raising the possibility that Dpp could signal between these cell types. However, in contrast to Hh ligand, Dpp is known to spread widely (Harmansa et al., 2015), even beyond the disc (Setiawan et al., 2018), and thus might regulate AMP gene expression in a more widespread manner. Individual plexins and semaphorins are differentially expressed in the epithelium and the AMPs, and could potentially mediate contact-dependent signaling between the two cell types. Finally, the receptor *robo2* is expressed in the indirect AMPs, while the gene that encodes its ligand *sli* was detected within a subset of AMPs (we observed higher levels of *sli* in direct AMPs, although it did not pass the criteria for significance by our differential expression analysis). The interaction of Sli with Robo causes repulsion of axonal growth cones at the midline during embryogenesis (Kidd et al., 1999). By analogy, the differential expression of Robo2 within the indirect AMPs, in conjunction with AMP expression of the Sli ligand, may serve to segregate the direct and indirect populations.

We have thus far not characterized a pathway where ligands secreted by the AMPs regulate gene expression in the epithelium. The gene encoding an EGF-family ligand *spitz* (*spi*) is expressed in the AMPs while its receptor is expressed in the epithelium. Also, two genes encoding poorly-characterized ligands, *miple1* and *miple2*, are expressed at especially high levels in the AMPs. Finally, the TNF ortholog *eiger* (Moreno et al., 2002; Kanda et al., 2002) is expressed by AMPs while the gene encoding its receptor *grindelwald* (*grnd*) (Andersen et al., 2015) is expressed in the disc epithelium. However, Eiger from the AMPs would not be expected to bind to Grnd since Grnd is expressed on the apical surface of disc cells. Transcripts for *wengen*, which encodes another potential Eiger receptor (Kanda et al., 2002), are expressed at high levels in the AMPs. Thus, our dataset provides many hints of signaling pathways that may function between the AMPs and the disc epithelium that provide multiple avenues for future investigations.

### Concluding remarks

Our work has provided a base for the study of heterotypic interactions in the developing wing disc during conditions of normal growth and demonstrate that such interactions can have a major effect on cell number, cell migration and cell fate in the wing disc. They also provide a useful starting point for investigations aimed at elucidating the role of heterotypic interactions under conditions of tissue damage and regeneration, overgrowth, or a wide variety of experimentally-induced perturbations.

## Materials and Methods

### Generation of single-cell suspension, barcoding, and sequencing

For each sample, approximately 250 staged *Drosophila* wing-imaginal discs were dissected within 1 hr. The collected tissue was then transferred to a microcentrifuge tube and incubated within a dissociation cocktail consisting of 2.5 mg/mL collagenase (Sigma #C9891) and 1X TrypLE (Thermo Fisher #A1217701) in Rinaldini solution (modified from Ariss et al., 2018). The sample tube was placed horizontally on a shaker machine operating at 225 rpm for 25 minutes at room temperature (method modified from Ariss et al., 2018). At the 10, 20, and 25 minute marks, the tube was flicked 20 times for additional mechanical dissociation. Dissociation was halted by centrifuging the sample at 5,000 rpm for 3 minutes, aspirating the dissociation cocktail, and then adding in 1 mL of cold PBS-10% FBS. The cell pellet was mixed by pipetting up-and-down approximately 25 times with a 1 mL pipette for additional mechanical dissociation, and then centrifuged again at 5,000 rpm for 3 minutes. The media was replaced with cold PBS-1% FBS, and the cell pellet was resuspended in preparation for FACS.

FACS of the sample was performed on a BD FACSAria Fusion flow cytometer. Dead cells were identified and removed via the addition of propidium iodide to the sample, and high-quality single cells were sorted into cold PBS-10% FBS. Cell concentration of the post-FACS sample was assessed by a hemocytometer, and adjusted 1,000 cells per uL.

Single-cell suspensions were barcoded for single-cell RNA sequencing with the 10X Chromium Single Cell platform (v2 chemistry). Barcoded samples were sequenced on an Illumina NovaSeq (S2 flow cell) to over 60% saturation.

### Single-cell data processing and analysis

The 10X Genomics Cell Ranger pipeline (v2.2.0) was used to align the sequencing reads to the *Drosophila melanogaster* transcriptome (version 6.24). The data was analyzed using the R and Python programming languages, primarily utilizing the packages Seurat v3 (Stuart et al., 2019) and scVI v0.4.1 (Lopez et al., 2018).

Our standard analysis pipeline is as follows: First, each dataset was analyzed separated using the standard Seurat pipeline, with no cells filtered, 30 principal components calculated, and clustering resolution set to 2.0 (all other parameters remained default). We then removed cell clusters with an abundance of low-quality cells (defined as clusters with mean number of genes detected per cell [nGene] was less than one standard deviation below the mean nGene of all cells in the dataset). Additionally, we found that each dataset had a cluster with markers for both AMP and epithelial cell types (e.g., *SPARC* and *Fas3*) and unusually high mean nGene; this cluster was suspected to be AMP-epithelial doublets, and was also removed. Clusters were then split into AMP and epithelial cell subsets based on the expression of known marker genes. Cells within each subset were subsequently filtered if either (1) their nGene count that was outside the mean nGene of the subset +/− 1.5 standard deviations, or (2) their percentage of reads for mitochondrial genes that was greater than 1.5 standard deviations above the mean mitochondrial read percentage of the subset.

Data subsets were harmonized into collective AMP or epithelium datasets using scVI. The scVI VAE model consisted of 2 layers (n_layers=2) and 20 latent dimensions (n_latent=20), with a negative-binomial reconstruction loss (reconstruction_loss=‘nb’). The model was trained on variable genes selected by Seurat’s variance-stabilizing transformation method; 1,000 (for epithelial subsets) or 2,000 (for AMP subsets) variable genes were calculated for each inputted batch, and then the union of these genes was supplied to scVI. The following parameters were used for model training: train_size=0.75, n_epochs=400, and lr=1e-3 (other parameters were left as default). Cell clustering and UMAP was performed using Seurat on the latent space derived from the scVI model. After harmonization, clusters were re-examined for doublet characteristics; clusters with a mean nGene count greater than one standard deviation above the mean nGene count of all cells were removed, as were clusters that displayed markers for both AMP and epithelial cell types. Identified hemocyte and tracheal cells were also separated out. scVI and Seurat were both re-run on the datasets to generate our final AMP and epithelial cell atlases (**Figures 2 and 3**).

To generate our full cell atlas consisting of all cell types (**Figure 1**), we merged and harmonized the cells in the AMP and epithelium cells atlases along with the separated hemocyte and tracheal cells. scVI and Seurat were run as previously described, with the scVI model trained on the union of the top 2,000 variable genes for each batch as calculated by Seurat. No additional cell filtering was performed after harmonization.

For visualizing data on UMAPs and dot plots, we calculated normalized and scaled expression counts using Seurat’s NormalizeData and ScaleData functions, respectively, with default parameters. For the normalized data, raw counts were normalized by total UMIs per cell, multiplied by 10,000. Natural-log normalized data is used for expression levels visualized with UMAP. For the scaled data, the natural-log normalized data is scaled for each gene, such that the mean expression is 0 with a standard deviation of 1. Scaled data is used for expression visualization on the dot plots.

### Cell sex and cell cycle correction with AMP data

Cells were classified as male or female by their expression levels of the dosage compensation complex genes *lncRNA:roX1* and *lncRNA:roX2* (Franke & Baker, 1999; Meller & Rattner, 2002), which are both expressed almost exclusively in male cells. For both genes, we examined the natural-log normalized expression counts (calculated by Seurat’s NormalizeData function), computed the density over the data, and identified the first local minima as a threshold (see **Figure 3 - figure supplement 1C**). Cells that were above the threshold for either *lncRNA:roX1* or *lncRNA:roX2* were classified as male; otherwise, they were classified as female. From this, we assigned 8,097 cells as male and 11,788 cells as female, which roughly matches the size ratio between male and female wing discs given that female discs are larger. We removed cell sex stratification by processing male and female AMPs as separate batches (for each actual batch) within scVI (**Figure 3 - figure supplement 1**).

We observed significant data stratification that correlated with a number of cell cycle-related genes, such as *Proliferating cell nuclear antigen* (*PCNA*) and *Cyclin B* (*CycB*) (Yamaguchi et al., 1990; Lehner & O’Farrell, 1990), indicating that our data was split between S phase and non-S phase (**Figure 3 - figure supplement 2**). Definitive classification of cells into cell cycle stages is difficult because expression of these genes is not typically demarcated sharply into specific cell cycle stages. To remove cell cycle stratification from our data, we examined the correlation of each scVI latent dimension with the expression levels of highly variable cell-cycle genes, and found that one latent dimension was strongly related (**Figure 3 - figure supplement 2D**). By masking this latent dimension from our downstream analysis (e.g., clustering and UMAP), we effectively diminished cell cycle stratification. **Figure 3A** shows the UMAP of our AMP data after subtraction of cell sex and cell cycle stratification, which allowed us to focus our analysis on different cell types within the AMPs (to see how each correction affected the AMP data, see **Figure 3 - figure supplement 2A-C**).

### Determining differentially-expressed genes

When examining clusters, genes were considered to have significant differential expression if they had (1) a false discovery rate < 0.05 (as calculated via Wilcoxon test), (2) a natural-log fold-change of 0.15 or more, and (3) a percent expression of at least 15% in one of the two populations in the comparison. This test was performed with Seurat’s FindMarkers function.

When evaluating differential expression between clusters of epithelial cells (e.g., **Figure 2H**), we used a one cluster vs. all analysis. When evaluating differential expression between direct and indirect AMPs (e.g., **Figure 3B**), we compared cells of the two groups as classified in **Figure 3C**. In these cases, differential expression statistics (i.e., FDR and fold change) are obtained by aggregating the cells (across batches) in each group and conducting a single comparison. When evaluating differential expression between time points (e.g., **Figures 1J, 2J**, and **3J**) (which would be inherently confounded with batch effects, since time points were collected across separate sequencing experiments), we took a conservative approach and only considered genes that were consistently significant (by the criteria defined above) in each temporal pairwise comparison (i.e., DE analysis was conducted between all temporal pairs: 96h1 vs. 120h1, 96h2 vs. 120h1, 96h2 vs.120h1, 96h2 vs.120h2). We report the natural-log of the average value for these pairwise comparisons, and the maximum FDR calculated (see **Supplementary files 1-3**).

### Generating a virtual model of the wing disc

We assembled reference gene expression patterns from a number of sources (Held Jr, 2002; Butler et al., 2003) and based our starting geometry on the disc proper from images in Bageritz et al., 2019. The images were processed in Adobe Photoshop and assembled in R with EBimage (Pau et al., 2010) to generate binarized gene expression reference for the AMPs, disc proper and peripodial epithelium. The geometry of the three-layered model is provided in **Supplementary file 4** and the binarized reference gene expression patterns are provided in **Supplementary file 5**. We used DistMap (Karaiskos et al., 2017) to statistically map single cells back to the reference. With this virtual wing disc model we used Distmap to calculate a ‘virtual *in situ’* or a prediction of gene expression patterns. This is based on the detected gene expression with the single-cell data and the mapping location to calculate relative expression values for our model. We mapped the AMP and epithelial cells separately, as this improved how well the model predicted genes with known expression patterns. In addition, we used the scVI imputed gene expression values when mapping the cells to the reference.

### Examination of receptor-ligand expression

From FlyBase, we assembled a list of genes encoding receptors and ligands from the following 19 pathways of interest: Wnt/Wingless, FGF, Hedgehog, PDGF/VEGF, JAK-STAT, Activin, BMP, Fat-Ds, Slit-Robo, Ephrin, Toll/Toll-Like, Semaphorin, Notch, Insulin-Like, Fog, Torso, Miple, EGFR, and TNF. For our analysis, we only examined pathways in which at least one receptor or ligand was either (1) differentially-expressed within one of the major domains of the epithelium (notum, hinge, pouch, or PE) when compared to all other epithelial cells, (2) differentially-expressed within one of the major domains of the AMPs (direct or indirect cells) when compared to each other, or (3) differentially-expressed within all epithelial cells or AMP cells when compared to each other. These pathways (and their receptors and ligands) are shown in **Figure 4H**.

### Drosophila stocks and husbandry

The stocks used in this study include the following lines: *R15B03-GAL4* (BDSC 49261); *G-TRACE* (BDSC 28280, 28281) (Evans et al., 2009); *pdm3-GFP* (BDSC 60560); *grn-GFP* (BDSC 58483); *ptc-GAL4* (BDSC *2017*); *dpp-GAL4* (BDSC 1553); *nub-GAL4; hh-GAL4; ap-GAL4; ci-GAL4*; *htl-GAL4* (*GMR93H07-GAL4,* BDSC 40669) is an enhancer within the first intron of the *htl gene. Drosophila* stocks from other labs: *UAS-ths* and *UAS-pyr* (A Stathopoulos). TRiP-CRISPR driven overexpression of *pyr* was conducted with a guideRNA that targets the upstream transcriptional start site, *P{TOE.GS00085}attP40* (BDSC 67537), and works together with a nuclease-dead Cas9 fused with a transcriptional activator domain, *UAS-dCas9.VPR* to cause gene activation (BDSC 67055) (Lin et al., 2015).

### Immunohistochemistry and image processing

Imaginal discs, unless otherwise noted, were fixed in 4% paraformaldehyde (PFA) for 15 min, permeabilized in PBS plus 0.1% Triton X-100 and blocked in 10% Normal Goat Serum. For anti-Nrt antibody staining, we permeabilized with (0.05%) Saponin. The following antibodies were used from the Developmental Studies Hybridoma Bank (DSHB): mouse anti-Cut (1:200, 2B10); mouse anti-Ptc (1:50, Apa-1); mouse anti-Nrt (BP 106 anti-Neurotactin); mouse anti-Wg (1:100, 4D4); mouse anti-En (1:10, 4D9); rat anti-Ci (1:10, 2A1); mouse anti-Ubx (1:20, FP3.38). The following antibodies were gifted: rat anti-Twist (1:1000, Eric Wieschaus), rabbit anti-Midline (1:500, James Skeath), and rat anti-Zfh2 (1:100, Chris Doe (Tran et al., 2010)). The following antibodies are from commercial sources: rabbit anti-DCP-1 (1:250, Cell signaling); rabbit anti-GFP (1:500, Torrey Pines Laboratories, Secaucus, NJ); chicken anti-GFP (1:500, ab13970 Abcam, Cambridge, UK); rabbit anti-beta-galactosidase (1:1000, #559762; MP Biomedicals, Santa Ana, CA). Secondary antibodies were from Cell Signaling. Nuclear staining with DAPI (1:1000).

Wing discs were imaged on a Zeiss Axioplan microscope with Apotome attachment, using 10x and 20x objectives. Image files were processed with ImageJ software. For each of the genotypes examined, we examined at least 8 discs and have reported representative results in this paper.

### Adult muscle preparations

To image adult flight muscles, male flies aged a minimum of 2 days after eclosion were anesthetized and submerged in 70% ethanol with dry ice. The thorax was isolated by removing the head, wings, legs, and abdomen. Thoraces were bisected sagittally with a 11-blade scalpel blade. For DVMs, the DLMs, leg muscles, and excess cuticle were removed from hemithoraces. For DFMs, the DLMs, DVMs, leg muscles, and excess cuticle were removed from hemithoraces. The DLMs, DVMs, and DFMs were fixed in 4% PFA for 2 hours. Muscles were then rinsed 3 times and permeabilized in 0.3% PBST for 3 cycles, 15 minutes each on a nutator. Hemithoraces were incubated in Rhodamine Phalloidin (1:200) and DAPI (1:500) in 0.3% PBST, then rinsed 3 times and washed in 0.3% PBST for 3 cycles, 15 minutes each on a nutator. Hemithoraces were mounted in a depression slide using antifade mountant. DLMs and DVMs were imaged with a 10x objective using a Zeiss Axioplan microscope. DFMs were imaged with the 20x and 63x objectives using confocal microscopy.

## Acknowledgments

The authors would like to thank A Stathopoulos, J Skeath, C Doe, and E Wieschaus for stocks and reagents and M Frolov for discussions. We thank current and former members of the Yosef lab for advice and guidance. We thank T Ashuach and D DeTomaso for discussions regarding computational analyses. We thank current and former members of the Hariharan lab for useful feedback and help with collection of samples. We thank C Wingert for technical assistance. We thank the Bloomington Stock Center, DRSC/TRiP Functional Genomics Resources, and Developmental Studies Hybridoma Bank for stocks and reagents. This work was funded by NIH grant R35 GM122490 and IKH is a Research Professor of the American Cancer Society (RP-16238-06-COUN).

---

**Figure 1 - figure supplement 1.**
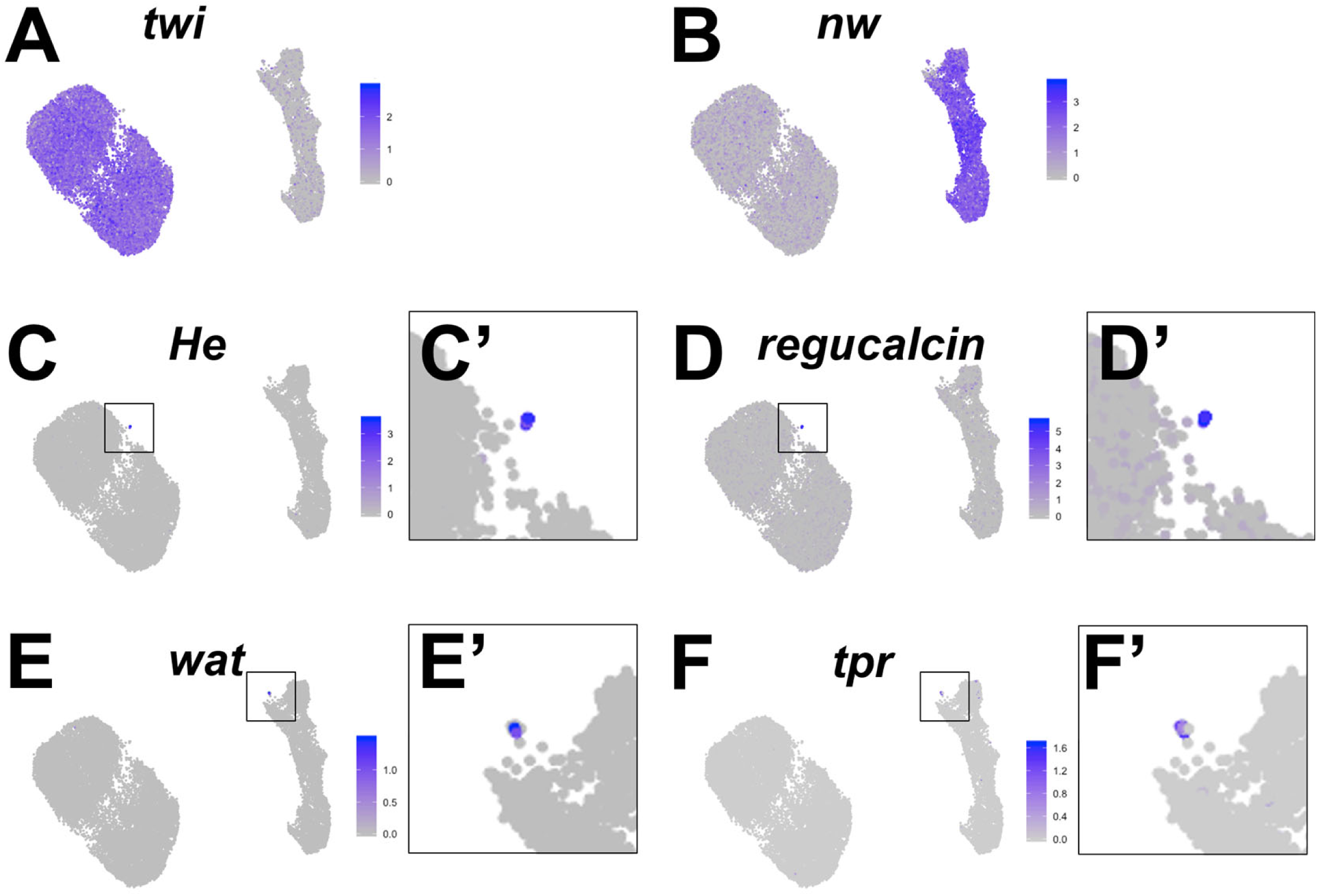
Cell type identification by known marker genes. UMAPs of full dataset showing expression levels for the following genes: AMP cell marker *twist (twi)* (**A**), epithelial cell marker *narrow wing (nw)* (**B**), hemocyte markers *Hemese* (*He)* (**C, C’**) and *regucalcin* (**D, D’**), and tracheal cell markers *waterproof* (*wat)* (**E, E’**) and *tracheal-prostasin (tpr*) (**F, F’**). Boxes in **C’**, **D’**, **E’** and **F’** are magnifications of indicated regions of the UMAP to show the cells that express the given marker gene. Color scales for UMAPs correspond to normalized (by total UMI) counts on a natural-log scale.

**Figure 2 - figure supplement 1.**
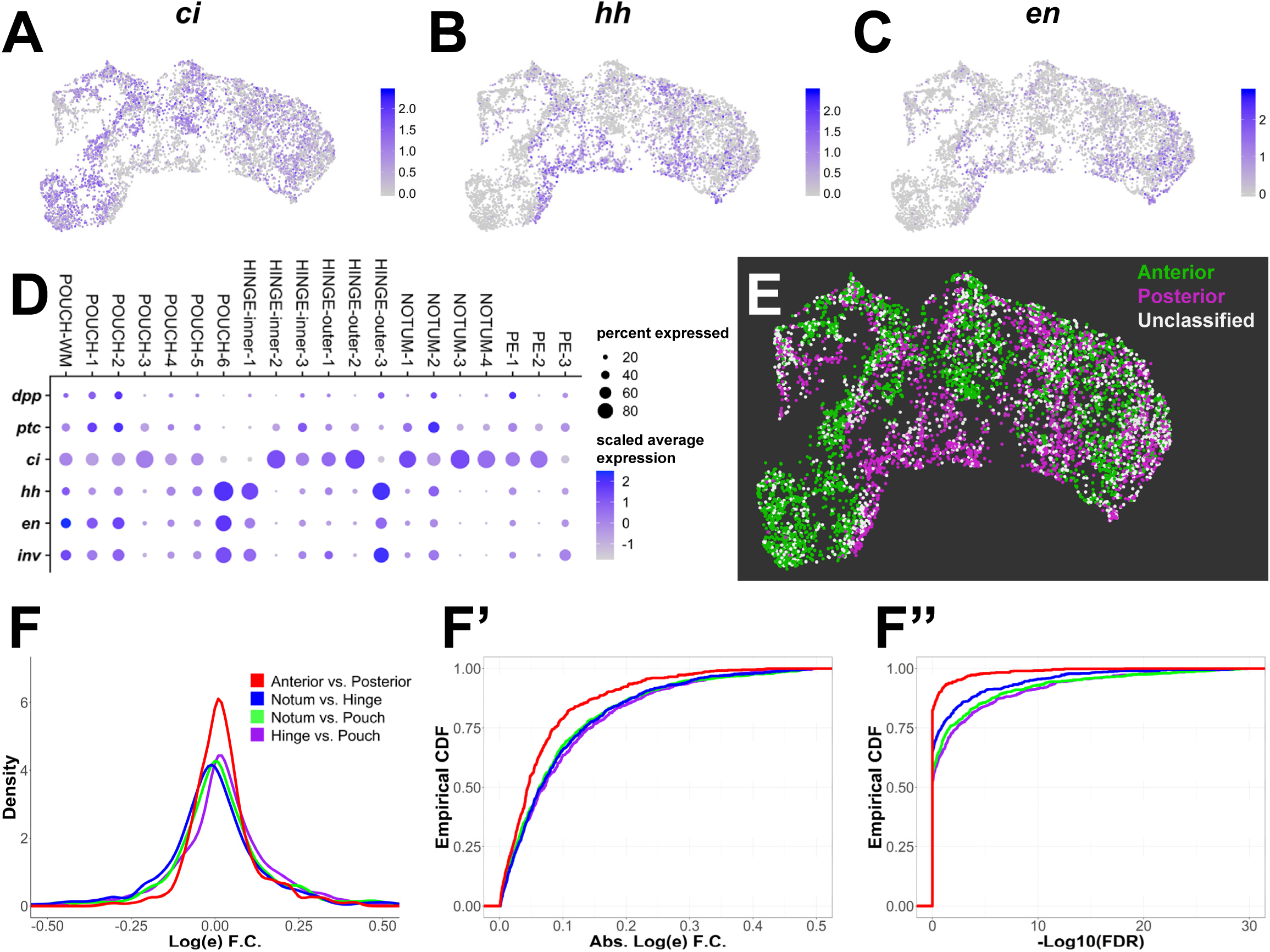
Anterior-posterior compartment identity is intermixed within the epithelium cell atlas. (**A-C**) UMAPs of disc epithelium single-cell data showing expression levels of anterior compartment marker *ci* (**A**) and posterior compartment markers *hh* (**B**) and *en* (**C**). (**D**) Dot plot summarizing expression of known anterior and posterior markers within the Seurat clusters of the disc epithelium data. Note that most clusters express both anterior and posterior markers, except for NOTUM-3, NOTUM-4, and HINGE-inner-2, which are primarily anterior, and POUCH-6 and HINGE-inner-1, which are primarily posterior (see also **Figure 4 - figure supplement 1**). (**E**) UMAP of anterior-posterior cell classification of disc epithelium data. Anterior cells have at least one transcript of anterior-compartment marker *ci*, whereas posterior cells have at least one transcript of posterior-compartment markers *hh*, *en*, or *inv*. Cells with transcripts for markers of both compartments or cells that lacked markers for either compartment were labeled as “Unclassified”. (**F**) Distributions of log fold-changes and FDR values of differential gene expression between regions of the disc epithelium. Differential expression was limited to variably-expressed genes within the disc epithelium. Panel **F** displays the density (or distribution) of log fold-changes of variable genes between regions of the epithelium. Panels **F’** and **F’’** empirical cumulative distribution function (ECDF) log fold-changes (**F’**) and FDR (**F’’**) of variable genes between regions of the epithelium. These ECDF plots are calculated as the percentage of variable genes (y-axis) below a particular log-fold change magnitude or FDR threshold. Note that differential gene expression between anterior and posterior cells is less dramatic than other comparisons; log fold-changes of variable genes between anterior and posterior cells are less extreme (the density is heavier around 0 in **F** and the ECDF has a steeper rise in **F’**), and the associated FDR values are less significant (**F’’**). Color scales for UMAPs correspond to normalized (by total UMI) counts on a natural-log scale.

**Figure 2 – figure supplement 2.**
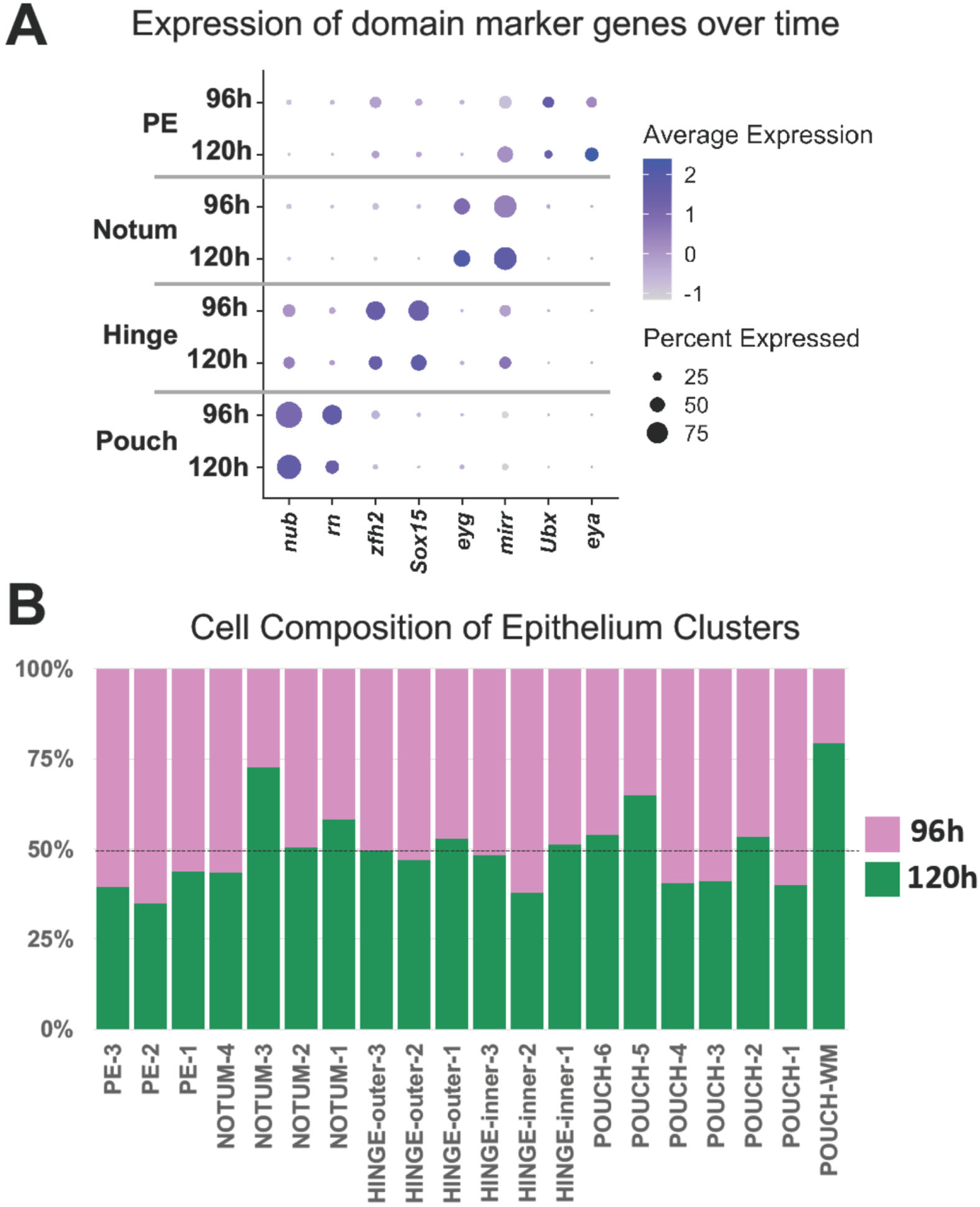
Markers of major epithelial domains and composition of cell clusters at developmental time points. (**A**) Dot plot showing the level and percent of marker gene expression for the major domains of the wing disc (pouch, hinge, notum, and peripodial epithelium (PE)) at both developmental time points, 96h and 120h. Note that the expression of these markers genes remains fairly constant over this developmental window. (**B**) Percent of cells in each of the epithelial cell clusters (as shown in **Figure 2G**) from the two developmental time points. Note that the “POUCH-WM” cluster has the highest percent of cells from the 120h time point and that “NOTUM-3” cluster is also enriched within the older sample. The other cell clusters are more evenly composed of cells from both developmental time points.

**Figure 3 – figure supplement 1.**
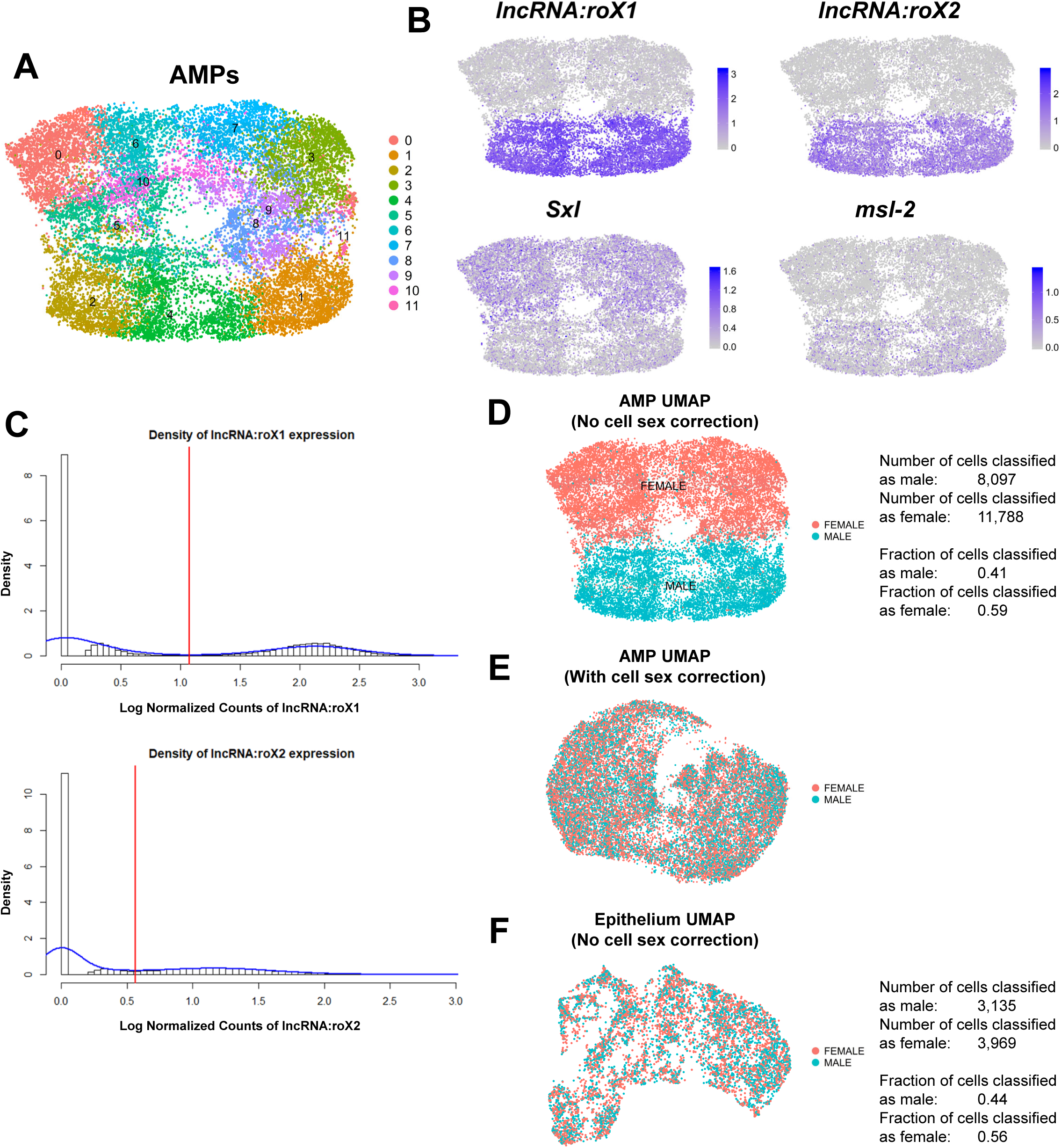
Correction of cell sex stratification within AMP scRNAseq data. (**A**) UMAP of AMP single-cell data, as processed by our standard computational pipeline. Colors correspond to computational cell clusters as determined by Seurat. (**B**) UMAPs showing expression of sex-specific genes *lncRNA:roX1*, *lncRNA:roX2*, *Sex lethal (Sxl)*, and *male-specific lethal 2 (msl-2)*. Note that the UMAPs show significant stratification based on the expression of these sex markers. (**C**) Probability histogram plots of the natural-log normalized expression counts for *lncRNA:roX1* and *lncRNA:roX2* within all cells. Density curves for the data are shown in blue. Red lines are drawn on the first local minima within the density of the data, and serve as a cutoff for classifying cells as having either high or low expression of either gene. Cells with high expression of either *lncRNA:roX1* or *lncRNA:roX2* were classified as male-originating; otherwise, cells were designated as female-originating. (**D**) Classification of cell sex shown on our standard-analysis UMAP of AMPs. Quantifications for the male-female classification are provided next to the UMAP. (**E**) Classification of cell sex shown via UMAP after computationally-correcting for cell sex stratification. Correction was performed by treating cell sex as a batch effect during data processing. Note that compared to panel **D**, male-originating and female-originated cells are now interspersed within the UMAP. (**F**) Classification of cell sex shown on our standard-analysis UMAP of epithelial cells. Quantifications for the male-female classification are provided next to the UMAP. Note that male-originating and female-originated cells are interspersed within the UMAP without need for correction. Color scales for UMAPs correspond to normalized (by total UMI) counts on a natural-log scale.

**Figure 3 – figure supplement 2.**
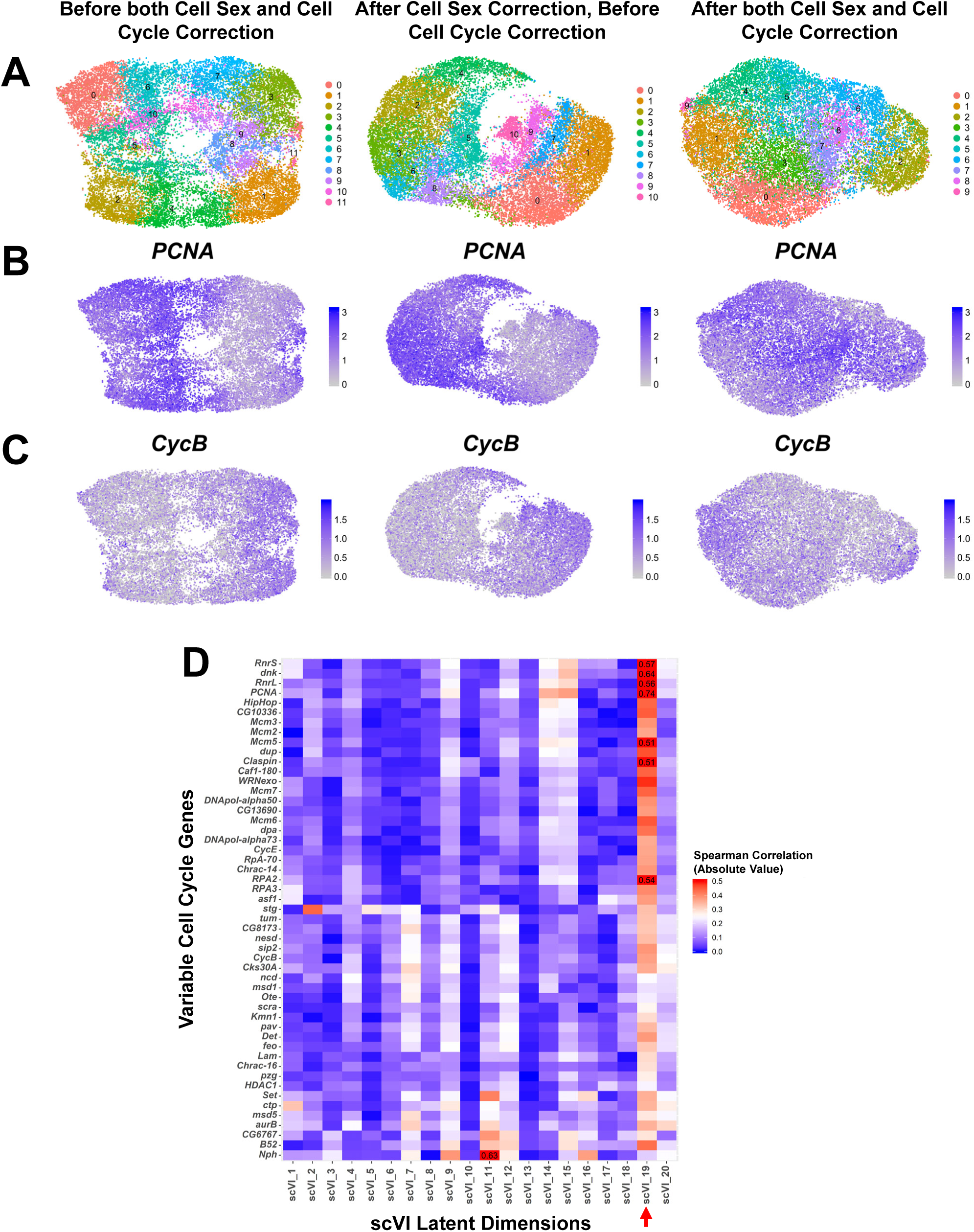
Correction of cell cycle stratification within AMP scRNAseq data. (**A-C**) UMAPs of AMP single-cell data without any data correction (first column), after cell sex correction only (second column) (see **Materials and Methods** and **Figure 3 – figure supplement 1** for details), and after both cell sex and cell cycle correction (third column). Cell cycle correction was performed by removing latent dimension scVI_19, which showed high correlation magnitude with variable cell cycle genes, from downstream analysis (e.g., data clustering and visualization) (see panel **D** of this figure and **Materials and Methods** for details). Cell colors correspond to Seurat clustering identities (**A**), or expression levels of cell-cycle genes *PCNA* (**B**) or *Cyclin B (CycB)* (**C**). Note that after we correct for cell cycle stratification, we observe better mixing of cell cycle markers *PCNA* and *CycB* throughout the data. (**D**) Magnitude of Spearman correlation between the scVI latent dimensions and the expression of variable cell cycle genes within the data. Color scale for correlation magnitude is capped at 0.5 for better visualization; correlation magnitudes that exceed 0.5 are written in the corresponding box. Note that scVI_19 (indicated on the x-axis by the red arrow) displays noticeably high correlation magnitude with most variable cell cycle genes, highlighting that it is capturing most cell cycle variation within the data. Color scales for UMAPs correspond to normalized (by total UMI) counts on a natural-log scale.

**Figure 3 – figure supplement 3.**
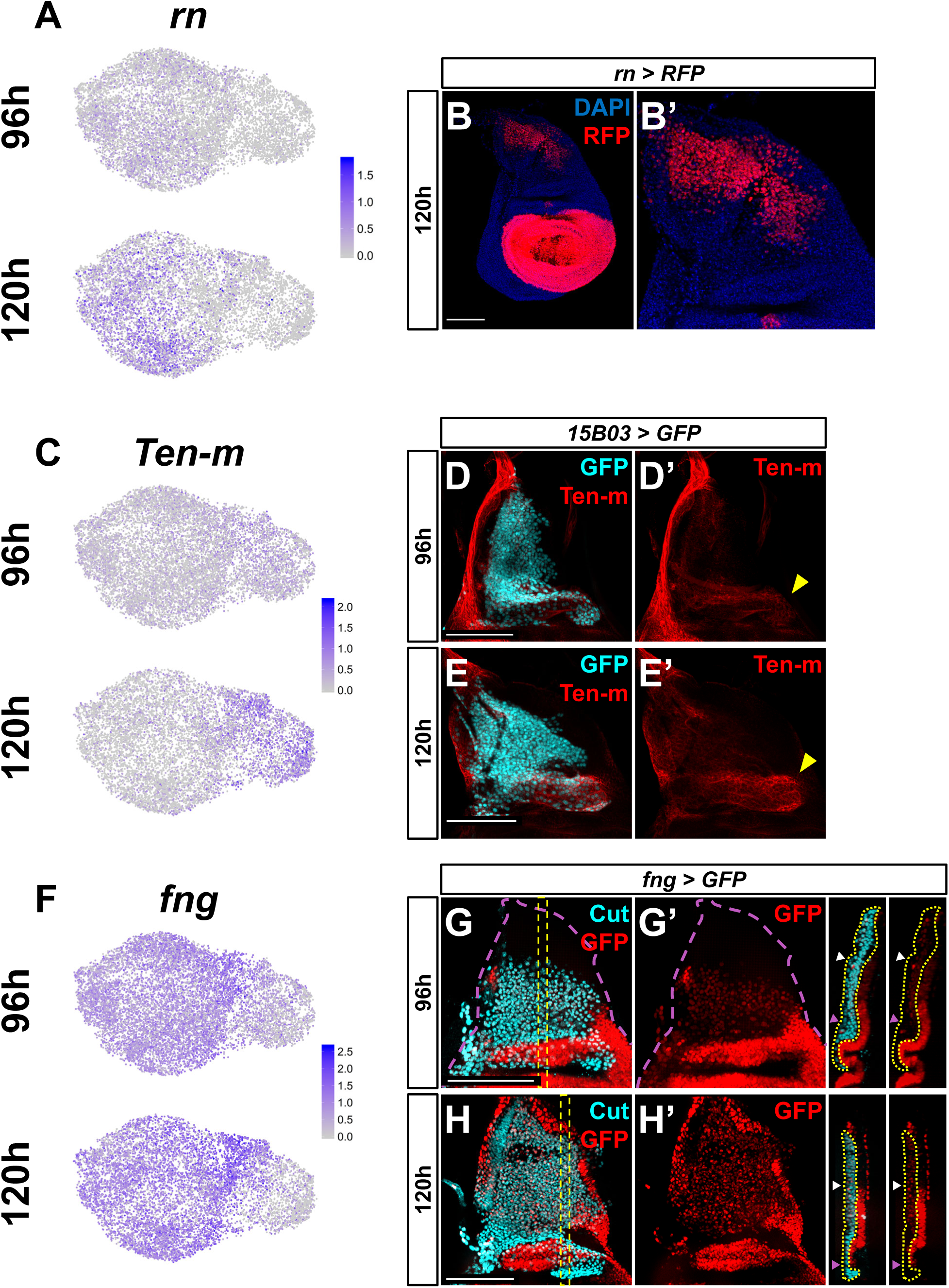
Gene expression patterns identified by AMP cell atlas. (**A, C, F**) UMAPs of *rn* (**A**), *Ten-m* (**C**), and *fng* (**F**) expression at 96h and 120h within AMPs. (**B**) Wing disc with RFP expression driven by *rn-Gal4*. AMPs shown at higher magnification in **B’**. (**D-E**) Wing discs stained for anti-Ten-m at 96h (**D, D’**) and 120h (**E, E’**). AMPs are visualized by expression of GFP via *15B03-Gal4* driver. Yellow arrowheads indicate regions of higher anti-Ten-m staining in direct AMPs. Note the increased levels of staining at 120h as compared to 96h. (**G-H**) Wing discs with GFP expression driven by *fng-Gal4* transgene. AMPs are visualized with anti-Cut. Magenta dashed lines in **G** and **G’** provide an outline of the wing disc. Orthogonal sections correspond to orthogonal max projections within the yellow dashed boxes in **G** and **H**. Yellow dotted lines within the orthogonal sections (apical is left, basal is right) outline the AMPs. White and magenta arrowheads indicate regions of high and low GFP fluorescence, respectively. Note the higher levels of GFP in more dorsal AMPs (white arrowheads) compared to more ventral AMPs (magenta arrowheads) at both time points. Color scales for UMAPs correspond to normalized (by total UMI) counts on a natural-log scale. Images shown in **B, D**, and **E** are max projections across image slices containing AMPs, whereas **G** and **H** are single slices through the AMPs. Microscopy scale bars = 100 μm.

**Figure 4 – figure supplement 1.**
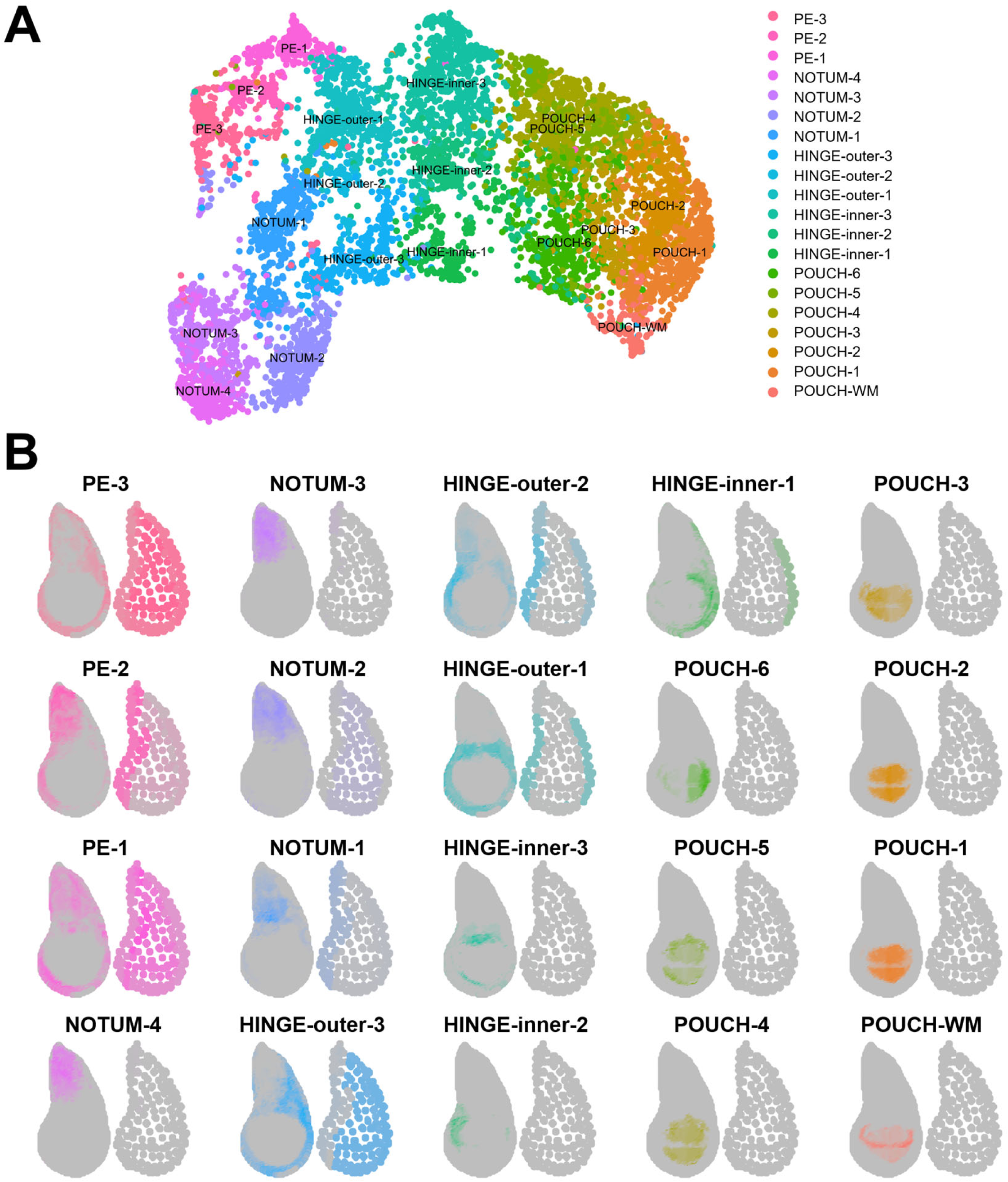
Mapping of epithelium cell clusters to the virtual wing disc. (**A**) UMAP of epithelium cell clusters (same as the UMAP in **Figure 2G**). (**B**) Visualization of where epithelium cell clusters map best to the reference model for the disc proper layer (on left) and the peripodial epithelium layer (on right). Gray regions indicate low predicted mapping, whereas regions with darker color shades indicate higher predicted mapping. The best cell mapping regions in (**B**) are colored to match the clusters in (**A**).

**Figure 4 – figure supplement 2.**
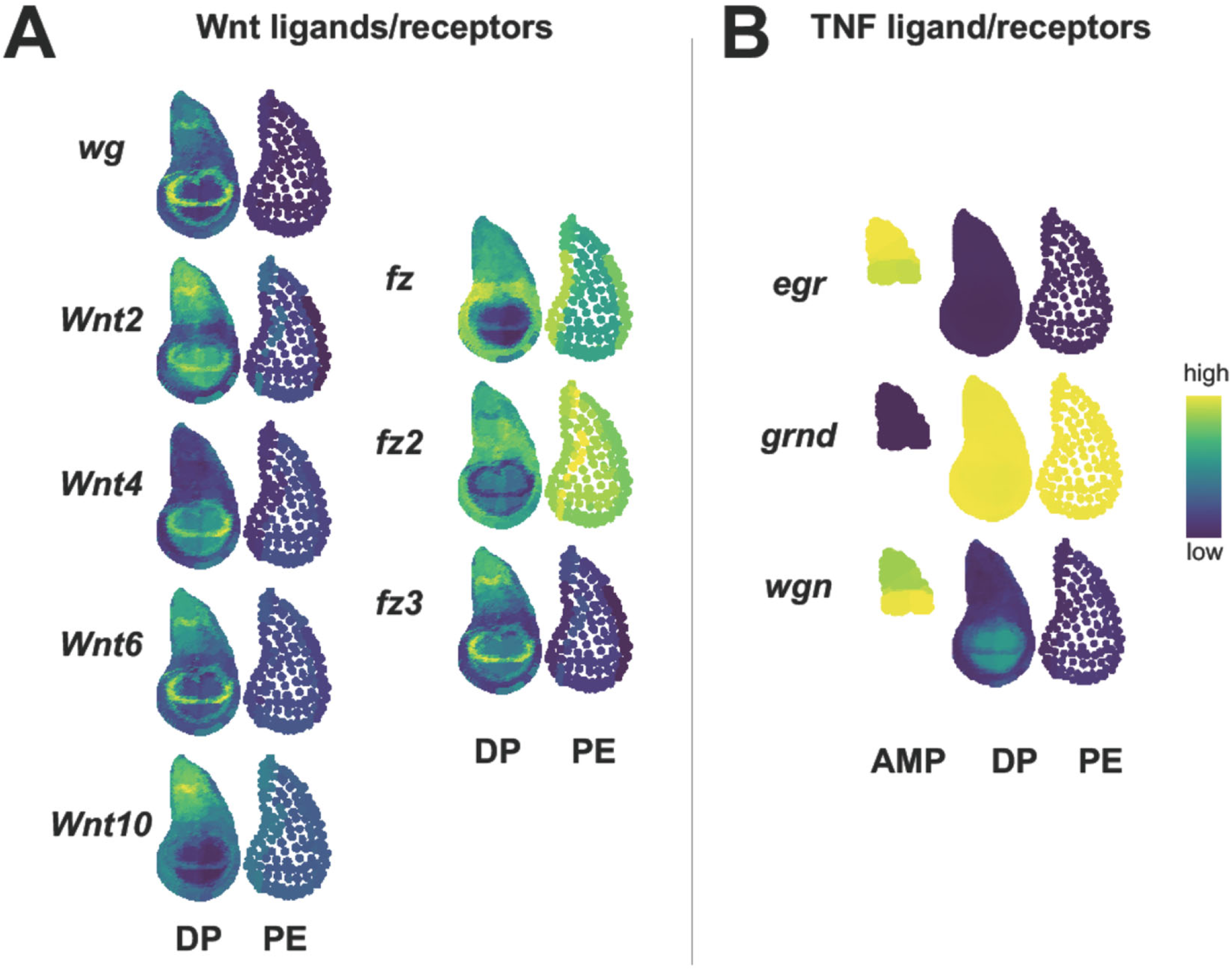
Predictions of gene expression for Wnt and TNF ligands and receptors. (**A, B**) Virtual *in situ,* or gene expression patterns predicted by our disc model. Scale bar on the right indicates the colors of relative predicted gene expression, with dark purple being low gene expression and yellow representing high expression. (**A**) Predicted gene expression patterns of the *Drosophila* Wnt ligands (*wg, Wnt2, Wnt4, Wnt6* and *Wnt10*) and *Drosophila* Wnt receptors (*fz, fz2,* and *fz3*) in the disc proper (DP) and peripodial epithelium (PE). (**B**) Predicted gene expression patterns for the *Drosophila* TNF ligand *(egr)* and its receptors (*grnd* and *wgn*) in the AMPs, disc proper (DP) and peripodial epithelium (PE). Note that *egr* is primarily expressed in the AMPs, while the receptor *grnd* is highly expressed within both layers of the epithelium.

**Figure 7 – figure supplement 1.**
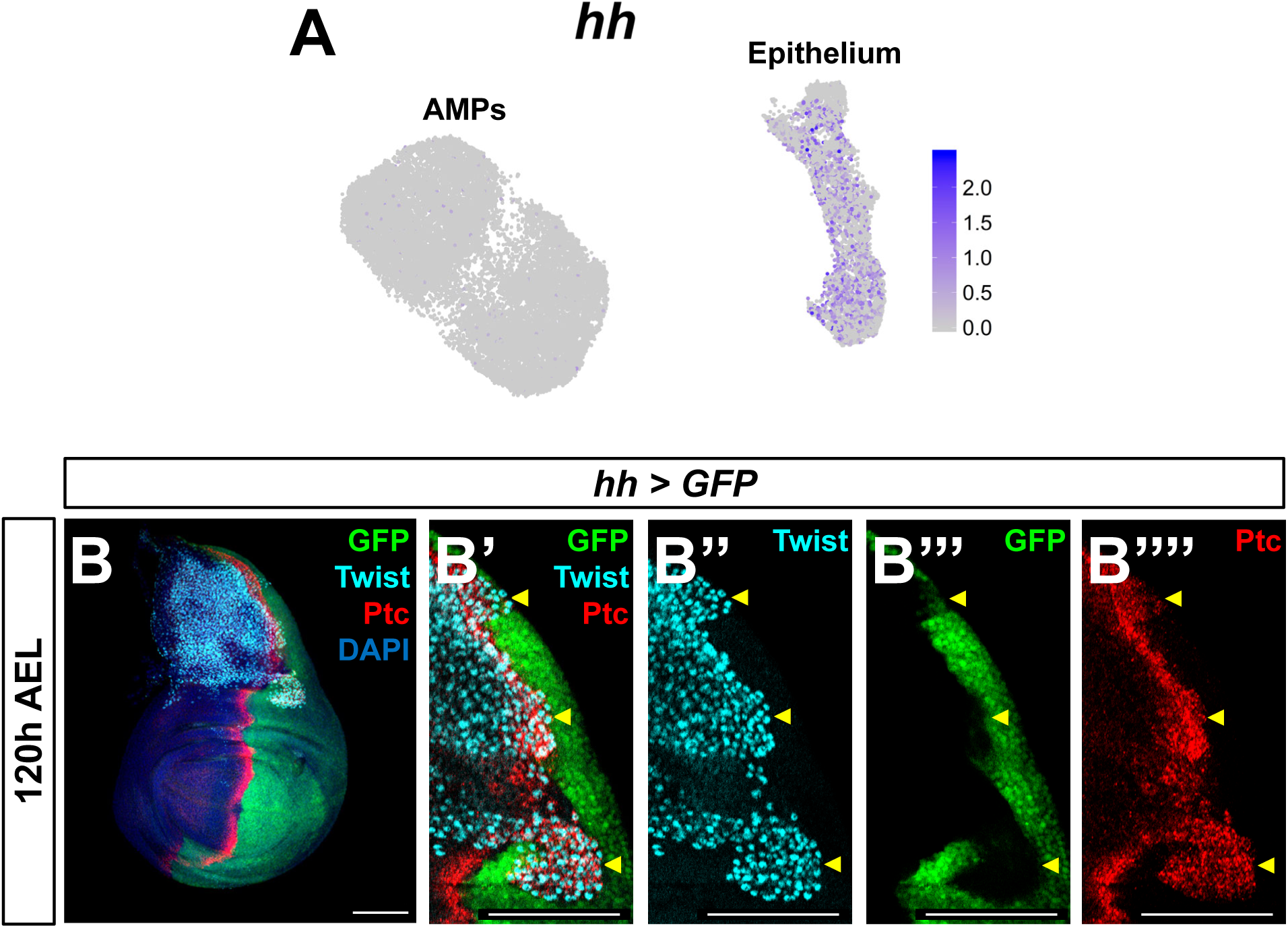
*hh* is expressed in the epithelium but not AMPs. (**A**) UMAP of disc epithelium and AMP cells (as in **Figure 1**), colored by expression of *hh*. Note the high levels of *hh* expression in the disc epithelium compared to the negligible expression within the AMPs. (**B**) Wing disc with GFP expression driven by *hh-Gal4* transgene. AMPs are visualized with anti-Twi staining. High-magnification images (**B’-B’’’’**) focus on posterior-localized AMPs, detectable by co-expression of anti-Twi and anti-Ptc staining and indicated by yellow arrowheads. Note that while the posterior compartment of the disc epithelium is labeled with GFP, the posterior-localized AMPs do not express *hh > GFP*. Color scales for UMAPs correspond to normalized (by total UMI) counts on a natural-log scale. Wing disc in **B** is a max projection over all image slices, whereas the high-magnification images in **B’-B’’’’** are single slices through AMPs. Microscopy scale bars = 100 μm.

**Figure 7 - figure supplement 2.**
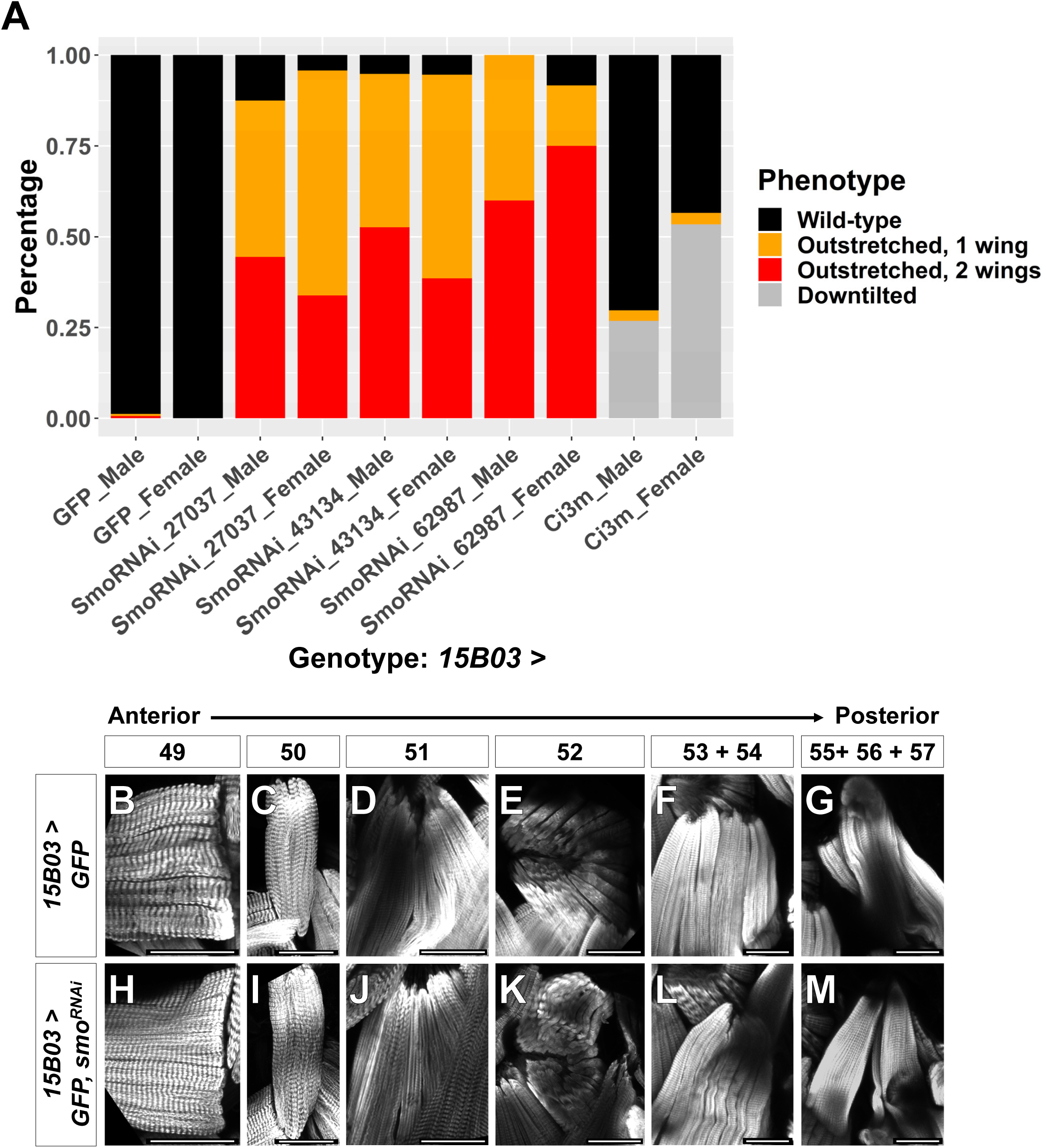
Adult wing posture phenotypes and morphology of individual muscle fibers after Hh-signaling perturbation. Quantification of observed wing posture phenotypes under conditions of Hh-signaling perturbation within AMPs (for description of wing posture phenotypes see **Figure 7**). All *UAS* lines are driven by *15B03-Gal4*. Data are presented for three different *UAS-smo^RNAi^* lines and separated by sex. Wing posture phenotypes were fairly consistent between males and females. Number of flies examined: GFP: 164 (male) and 179 (female), *smo^RNAi^* Bloom. #27037: 72 (male) and 71 (female), *smo^RNAi^* Bloom. #43134: 135 (male) and 148 (female), *smo^RNAi^* Bloom. #62987: 20 (male) and 12 (female), *ci^3m^*: 175 (male) and 191 (female) (**B-O**) Major DFMs in adults flies where *15B03-Gal4* drives *>GFP* alone (**B-H**) or *>GFP* together with *>smo^RNAi^* (**I-O**). Muscle fibers are shown in the order of relative anterior-posterior positioning within the thorax, with numbering nomenclature as described in **Figure 7O**. Note that in *>smo^RNAi^* flies, the posterior edge of muscle 52 appears to be truncated (**L**), muscles 53 and 54 are indistinguishable and both project to the dorsal attachment site of muscle 54 **(M, N, Figure 7U)**, and muscles 55, 56, and 57 are disorganized **(O)**. Microscopy scale bars = 50 μm.

**Figure 8 – figure supplement 1.**
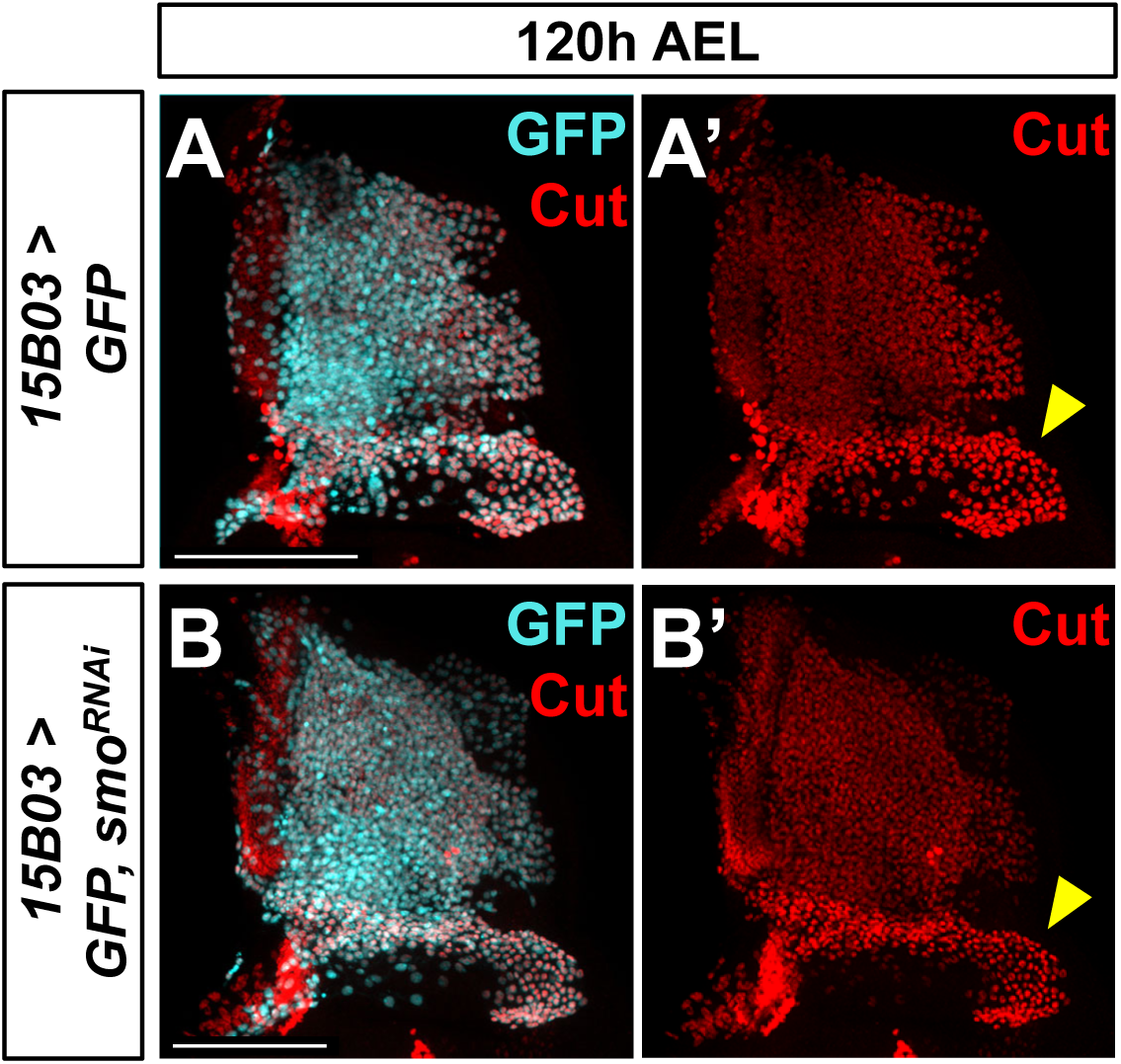
Knockdown of *smo* does not affect Ct protein levels in AMPs. (**A, B**) Close up of the notum region of wing discs stained for anti-Ct (red) with AMP-specific *15B03-Gal4* driving *>GFP* alone (**A**) or *>GFP* with *>smo^RNAi^* (**B**). Yellow arrowheads indicate higher levels of Ct staining in direct AMPs. Note that Ct staining in direct AMPs is unaffected by *smo^RNAi^* expression. Microscopy scale bars = 100 μm.

**Supplementary file 1. Genes with differential expression between 96h and 120h within the epithelium and AMPs.** Genes were selected based on being significantly and consistently upregulated or downregulated between the two time points in either the disc epithelium and/or the AMPs. The average gene expression within cells (natural-log scale), fraction of cells expressing a given gene, fold-change between time points (natural-log scale), and FDR for differential-expression significance are reported. These gene expression, detection, and fold-change calculations are averaged across each of the pairwise comparisons performed, and the max FDR value is shown (see **Materials and Methods** for details on differential expression between time points). Negative fold-change values indicate higher expression at 96h and are colored magenta. Positive fold-change values indicate higher expression at 120h and are colored green. N.R. = not replicable; calculations in which the fold-change direction differed between pairwise comparisons.

**Supplementary file 2. Genes with differential expression between 96h and 120h within the epithelial cell clusters.** Genes were selected based on being significantly and consistently upregulated or downregulated between the two time points in at least one epithelial cluster. The natural-log of the fold change between 96h and 120h is reported, averaged across each of the pairwise comparisons performed (see **Materials and Methods** for details on differential expression between time points). Negative values indicate higher expression at 96h and are colored magenta. Positive values indicate higher expression at 120h and are colored green. Values that were not significant (based on max FDR) are reported with a “-”.

**Supplementary file 3. Genes with differential expression between 96h and 120h within the direct and indirect AMP.** Genes were selected based on being significantly and consistently upregulated or downregulated between the two time points in either the direct and/or the indirect AMPs. The average gene expression within cells (natural-log scale), fraction of cells expressing a given gene, fold-change between time points (natural-log scale), and FDR for differential-expression significance are reported. These gene expression, detection, and fold-change calculations are averaged across each of the pairwise comparisons performed, and the max FDR value is shown (see **Materials and Methods** for details on differential expression between time points). Negative fold-change values indicate higher expression at 96h and are colored magenta. Positive fold-change values indicate higher expression at 120h and are colored green. N.R. = not replicable; calculations in which the fold-change direction differed between pairwise comparisons; calculations in which the fold-change direction differed between pairwise comparisons.

**Supplementary file 4. Geometry of disc model.** CSV file of the X, Y, Z geometry used in reference gene expression patterns (**Supplementary file 5**). Formatted as used in DistMap to generate virtual wing-disc.

**Supplementary file 5. Reference gene expression patterns.** CSV file of the binarized reference gene expression patterns (along with geometry in **Supplementary file 4**). Formatted as used in DistMap to generate virtual wing-disc.

